# antiCD49d Ab treatment ameliorates age-associated inflammatory response and mitigates CD8+ T-cell cytotoxicity after traumatic brain injury

**DOI:** 10.1101/2024.06.17.596673

**Authors:** Zhangying Chen, Kacie P. Ford, Mecca B.A.R Islam, Hanxiao Wan, Hyebin Han, Abhirami Ramakrishnan, Ryan J. Brown, Veronica Villanueva, Yidan Wang, Booker T. Davis, Craig Weiss, Weiguo Cui, David Gate, Steven J. Schwulst

## Abstract

Patients aged 65 years and older account for an increasing proportion of patients with traumatic brain injury (TBI). Older TBI patients experience increased morbidity and mortality compared to their younger counterparts. Our prior data demonstrated that by blocking α4 integrin, anti-CD49d antibody (aCD49d Ab) abrogates CD8+ T-cell infiltration into the injured brain, improves survival, and attenuates neurocognitive deficits. Here, we aimed to uncover how aCD49d Ab treatment alters local cellular responses in the aged mouse brain. Consequently, mice incur age-associated toxic cytokine and chemokine responses long-term post-TBI. aCD49d Ab attenuates this response along with a T helper (Th)1/Th17 immunological shift and remediation of overall CD8+ T cell cytotoxicity. Furthermore, aCD49d Ab reduces CD8+ T cells exhibiting higher effector status, leading to reduced clonal expansion in aged, but not young, mouse brains with chronic TBI. Together, aCD49d Ab is a promising therapeutic strategy for treating TBI in the older people.

**Graphic abstract:** 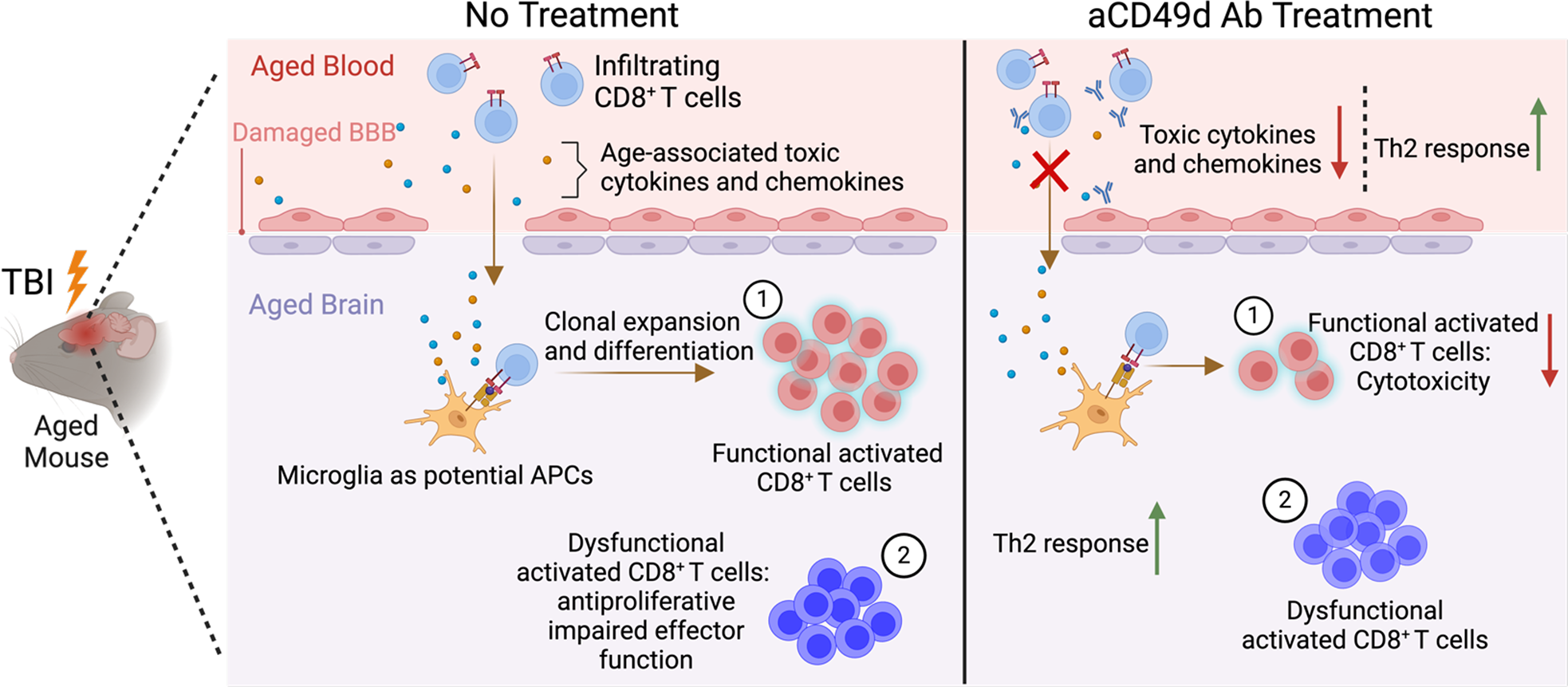

**Aged brains after TBI comprise two pools of CD8^+^ T cells**. The aged brain has long been resided by a population of CD8^+^ T cells that’s exhaustive and dysfunctional. Post TBI, due to BBB impairment, functional CD8^+^ T cells primarily migrate into the brain parenchyma. Aged, injury-associated microglia with upregulated MHC class I molecules can present neoantigens such as neuronal and/or myelin debris in the injured brains to functional CD8+ T, resulting in downstream CD8+ T cell cytotoxicity. aCD49d Ab treatment exerts its function by blocking the migration of functional effector CD8^+^ T cell population, leading to less cytotoxicity and resulting in improved TBI outcomes in aged mice.

## Introduction

Adults aged 65 years and older account for over 40% of all TBI-related hospitalizations in the United States (1, 2). In addition, aged TBI patients experience higher morbidity, slower recovery, and worse outcomes as compared to younger TBI patients (3). Hence, TBI in older adults is an underrecognized public health concern. Yet, age-specific treatment paradigms are lacking. To overcome this challenge, it is paramount to identify age-specific mechanisms for the worse outcomes observed in aged TBI patients. With normal aging, there is an infiltration of CD8^+^ T into the brain parenchyma, likely due to recruitment signals from aged microglia as well as age-related permeability of the blood-brain barrier (BBB) (4–8). As previously published by our group, CD8^+^ T cell infiltration is further increased after TBI leading to a substantial accumulation of CD8^+^ T cells within the brains of aged subjects. Consequently, CD8^+^ T-cell infiltrates can directly contribute to neuroinflammation and cognitive impairment in normal aging, injury, and numerous neurodegenerative diseases (7–8).

Previous work in our laboratory demonstrated that blocking CD8^+^ T cell infiltration with the FDA-approved drug, Natalizumab, is highly effective in improving TBI outcomes in aged mice. Natalizumab is a monoclonal antibody against integrin α4 (aCD49d Ab) of leukocytes. It is a current FDA-approved treatment for multiple sclerosis (MS) and works by preventing peripheral lymphocyte extravasation into the brain and spinal cord, thereby reducing inflammation and nerve damage (9). In our previously published work, we found that aCD49d Ab significantly reduced the infiltration of CD8^+^ T cells into the injured brain, improved survival after TBI, and ameliorated neurocognitive and motor deficits in our murine model of aged TBI whereas no therapeutic effect was observed in young mice after TBI (8). In the periphery, repeated aCD49d Ab treatment augmented the Th2 response, indicated by significantly increased levels of the anti-inflammatory cytokines IL-4 and IL-13 within the plasma of aged TBI mice (8). However, in the setting of TBI, the molecular mechanisms underlying the therapeutic effects of aCD49d Ab have yet to be determined.

In addition to lymphocyte migration into the CNS, the brain is home to a specialized resident immune cell component. While parenchymal microglia and their functions have been the subject of much scrutiny over the years, the non-parenchymal monocyte-derived macrophages, known as border-associated macrophages (BAMs), are only recently recognized as having a significant role in injury and disease. BAMs reside within the dura mater, subdural space, and choroid plexus (CP) which contain fenestrated blood vessels forming the interface with the peripheral circulation (10). With aging and disease states, a portion of the overall BAM population is replaced by monocytes from the periphery (i.e., monocyte-derived macrophages) (10–12). Thereby, we surmised aCD49d Ab treatment would also affect brain macrophage dynamics through modulating peripheral immunity and cytokine and chemokine response. Taken together, the current study is focused on the molecular mechanisms of aCD49d T cell blockade in TBI. We performed an unbiased analysis via single-cell RNA sequencing coupled with T cell receptor (TCR) sequencing of CD45+ immune cells isolated from the brains of young and aged mice at 2 months post TBI. Molecular changes were further integrated with high-parameter flow cytometry and multiplex cytokine assays to systemically study how repeated aCD49d Ab treatment alters local downstream cellular responses within the aged mouse brain.

## Results

### aCD49d Ab Treatment Reduced CD8+ T cells within Aged Brains and Improved Deficits in Motor Function and Working Memory After TBI

TBI was induced in mice via controlled cortical impact (CCI) as we have previously published (7,8). All mice underwent a 1-cm sagittal incision to expose the skull. Sham mice only received the incision whereas TBI mice further received a 5-mm craniectomy to expose the brain at coordinates, −2 mm anteroposterior axis and +2 mm mediolateral of bregma. Following craniotomy, TBI mice received a severe injury at 2-mm depth delivered by a 3-mm impacting tip striking the exposed cortex at 2.5 m/s. At a clinically relevant time of 2 hours post TBI or sham surgery, mice received 300 ug of aCD49d AB or its isotype control via intraperitoneal injection. Dosing was repeated every two weeks for up to 2 months. Brains were harvested at 2 months post TBI. As expected, in the aged mouse brains 2 months post TBI, aCD49d Ab treatment led to an attenuation of lymphocyte accumulation (Supp Fig. 1Ai and ii) and specifically reduced CD8^+^ T cells (Supp Fig. 1Aiv) but not CD4^+^ T cells (Supp Fig. 1Aiii) in the brain parenchyma compared to isotype treatment. Meanwhile, the CD8^+^ T cell reduction was not noted within the brains of young mice (Supp Fig. 1Aiv, 8). Furthermore, to characterize the effect of aCD49d Ab treatment on motor coordination and spatial working memory, deficits in which are common in human TBI patients, mice underwent rotarod and Y-maze tests. In the rotarod test, aged aCD49d Ab treated TBI mice spent more time on the accelerating wheels, indicative of improved motor coordination as compared to isotype treated mice (Fig. 1A). Analysis of spatial working memory is assessed by recording the spontaneous alternations between the two arms of a Y-shaped maze. Over the course of the test, a naïve mouse tends to enter the less recently visited arm. In the current study, aged aCD49d Ab treated TBI mice had a higher percent alternation score than isotype treated TBI mice (Fig. 1B). In fact, the aged aCD49d Ab treated mice demonstrated similar performance to aged sham injured mice indicating preservation of working memory after injury. In the meantime, consistent to prior findings, no effect of aCD49d Ab was seen in young mice post TBI (Supp Fig. 2).

**Figure 1.**
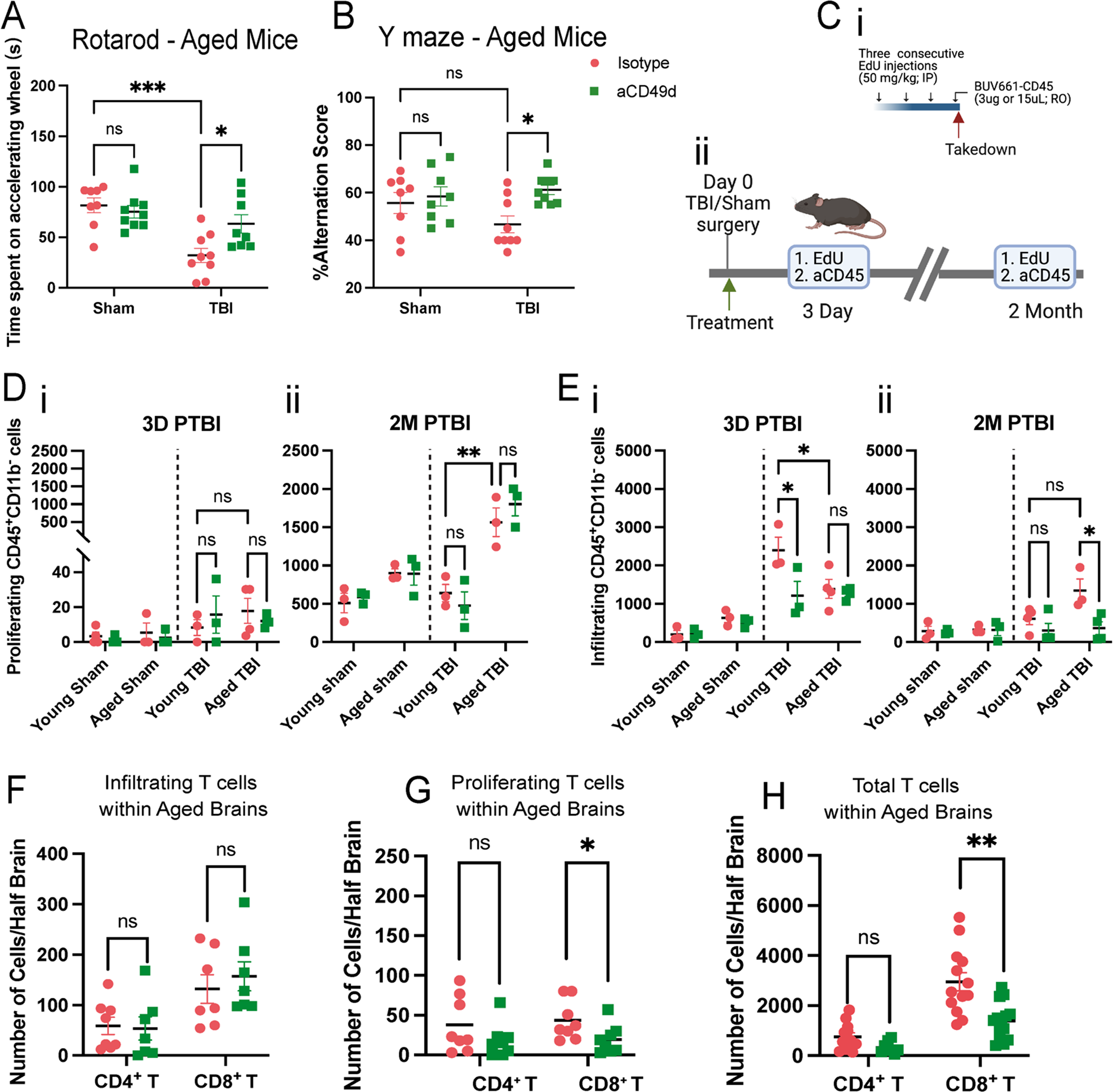
aCD49d Ab treatment reduced the infiltration and total number of CD8^+^ T cells in the injured brains of aged mice. A. Results of rotarod indicated by time spent on accelerating wheels (s). B. Results of Y maze indicated by %alteration score. C. i. A scheme of the EdU and BUV661-CD56 injections. IP, intraperitoneal injection. RO, retro-orbital injection. ii. Mice received EdU and BUV661-CD45 to label proliferating and infiltrating cells in the brains at 3 days and 2 months post-TBI. D. Quantifications of infiltrating CD45^+^ cells in mouse brains at i. 3 days post-TBI and ii. 2 months post-TBI. E. Quantifications of proliferating lymphocytes in mouse brains at i. 3 days post-TBI and ii. 2 months post-TBI. Quantifications of F. infiltrating, G. proliferating, H. total T cells. Data are from one independent experiment in D-E and two independent experiments in A-B and F-H. All data are shown as the mean ± SEM. 2-way ANOVA with Tukey’s multiple comparisons test in A-B and D-E, n = 8-10/group for A and B, n = 3-4/group for D and E,.Student’s t-test in F-H, n = 7-13/group. *p < 0.05, **p < 0.01, ***p < 0.001.

To assess CD49d expression in the periphery among different immune cell types at a steady state, we sampled 25 μL of blood via submandibular collection from B6 naïve mice of different ages. Remarkably, both CD4+ and CD8+ T cells exhibit age-dependent upregulation of CD49d expression (Supp Fig. 3), indicating an increased potential of T cell extravasation and migration with aging.

### aCD49d Ab Treatment Decreased Lymphocyte Infiltration and Reduced the Number of CD8^+^ T cells in the Injured Brains of Aged Mice

To distinguish local proliferation versus infiltration of T cells in post-TBI brains, 5-ethynyl-2’-deoxyuridine (EdU), a well-established thymidine analogue assaying DNA synthesis, was administered intraperitoneally for three consecutive days and BV611 conjugated anti CD45 antibodies were injected retro-orbitally two hours prior to euthanasia in order to label proliferating and infiltrating cells, respectively (Fig. 1Ci and ii). Intracardiac perfusion was performed to clear the vasculature. Brains were harvested which were used to compare the rate of infiltrating versus proliferating cells at 3 days (acute) and 2 months (chronic) post-TBI (gating strategy for CD45^+^ and EdU^+^ brain cells shown in Supp Fig. 1B and C). At 3 days post-TBI, young mice had significantly more infiltrating CD45^+^ cells than aged mice (Fig. 1D). However, at 2 months post-TBI, aged mice had more infiltrating CD45^+^ cells than young mice (Fig. 1D). As expected, aCD49d Ab limited the infiltration of CD45^+^ cells (Fig. 1D), especially in aged mice 2 months post-TBI. Focusing on local proliferating cells, we observed only few proliferating CD45^+^ cells at 3 days post-TBI (Fig. 1E). Contrastingly, we noted increasing proliferating CD45^+^ cells, more prominent in aged than young brains 2 months post-TBI (Fig. 1E). Phenotypic analysis of T cells revealed that aCD49d Ab treatment significantly reduced infiltrating CD8^+^ T cells (Fig. 1F) but had no effect on local proliferating T cells (Fig. 1G) in aged TBI brains. Hence, total CD8^+^ T within TBI brains were also significantly reduced in the aged aCD49d Ab-treatment group as compared to the isotype-treated group (Fig. 1H).

### aCD49d Ab Treatment Suppressed Peripheral and Local Inflammatory Immune Responses in Acute TBI

Next, a Luminex multiple cytokine assay was performed to assess cytokine and chemokine levels in the blood and within the brain. Sham mice, regardless of treatment, had no above-threshold detection of injury-related cytokines and chemokines. Therefore, their data were shown as zero in Fig. 2. Consequently, plasma levels of the pro-inflammatory cytokines interlukin-6 (IL-6), IL-17A, and IL-22 as well as tumor necrosis factor alpha (TNF-α) were elevated in aged TBI mice 3 days post-injury as compared to young mice (Fig. 2Ai-iii and v). However, aCD49d Ab treatment markedly reduced their levels in aged TBI mice (Fig. 2Ai-iii and v). Similar suppression after aCD49d Ab treatment was also noted in CXC chemokine ligand 2 (CXCL2), which is produced almost exclusively by neutrophils, in aged TBI mice at 3 days post-injury as compared to isotype treated mice (Fig. 2Aiv). Furthermore, IL-12p70 (IL-12), which is produced by activated antigen-presenting cells (APC) and enhance Th1 and cytotoxic CD8+ T cells, along with IL-1β were elevated at 7 days post-TBI in isotype treated mice and were decreased with aCD49d Ab treatment (Supp Fig. 4A and B). Interestingly, the peripheral cytokine response mirrored that in the brain as observed in IL-6, IL22, IL-17A, and CXCL2 (Fig. 2B i-iv). There were no significant changes, or expression was below the level of detection, in the remaining cytokines and chemokines analyzed in any group.

**Figure 2.**
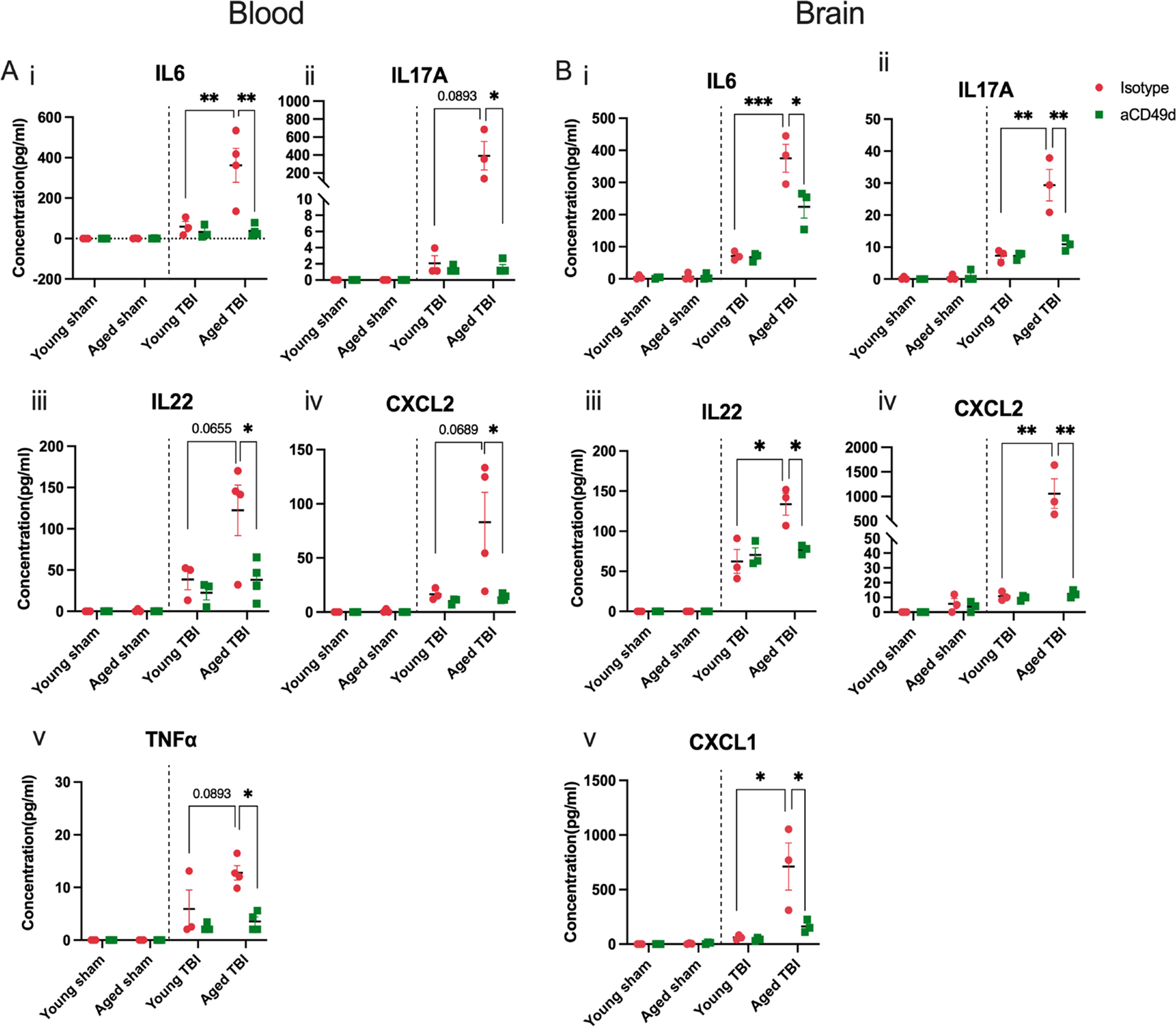
Multiplex cytokine analysis in aged and young mice at 3 days post injury. A. Levels of cytokines in the plasma including i. CXCL2, ii. IL-22, iii. IL-6, iv. IL-17A, and v. TNF-α. B. Levels of cytokines in the ipsilateral brain tissue homogenates including i. CXCL1, ii. CXCL2, iii. IL-6, iv. IL-22, and v. IL-17A. Data are shown as the mean ± SEM, 2-way ANOVA with Tukey’s multiple comparisons test. n = 3-4/group, *p < 0.05, **p < 0.01, *** p < 0.001. Data are from one independent experiment.

### aCD49d Ab Treatment Suppressed Immune Cell Chemotaxis but Induced GM-CSF in Aged Brains with Chronic TBI

At 2 months post TBI, several chemoattractants continued to increase within aged mouse brains compared to young mouse brains. aCD49d Ab treatment suppressed the level of CXCL1 and CCL11 which are involved in the selective recruitment of neutrophils and eosinophils, respectively (Fig. 3Ai and iii). Likewise, CCL7, a potent chemoattractant that recruits multiple leukocyte populations, showed similar suppression with aCD49d Ab treatment (Fig. 3Aii). To our surprise, granulocyte-macrophage colony-stimulating factor (GM-CSF) in the aged mouse brains was comparable to the level in young mouse brains 2 months post TBI. However, GM-CSF levels were found to be increased with aCD49d Ab treatment along with increases in CCL2 and TNF-α (Fig. 3Aiv-vi). To measure cytokine expression capacity within the brains isolated from aged TBI mice, we stimulated brain derived immune cells with PMA/ionomycin and incubated with golgi-plug containing brefeldin A *ex vivo* (Fig. 3B). Consequently, CD4+ T cells expressing IL-13 were significantly increased in the aCD49d Ab-treated brain cells compared to the isotype-treated brain cells isolated from aged TBI mice (Fig. 3B). CD4+ T cells express transcription factors (TF) T-bet, GATA-3, RORγt, representing Th1, Th2, and Th17. As a result, aged mouse brains had decreased frequency of T-bet^+^, GATA-3^+^ yet elevated frequency of RORγt^+^ CD4+ T cells compared to young mouse brains (Fig. 3C). aCD49d Ab appeared to not affect the frequency of CD4+ T cell subtypes except a further decrease in the frequency of T-bet^+^ in CD4+ T cells in aged mice 2 months post TBI (Fig. 3C).

**Figure 3.**
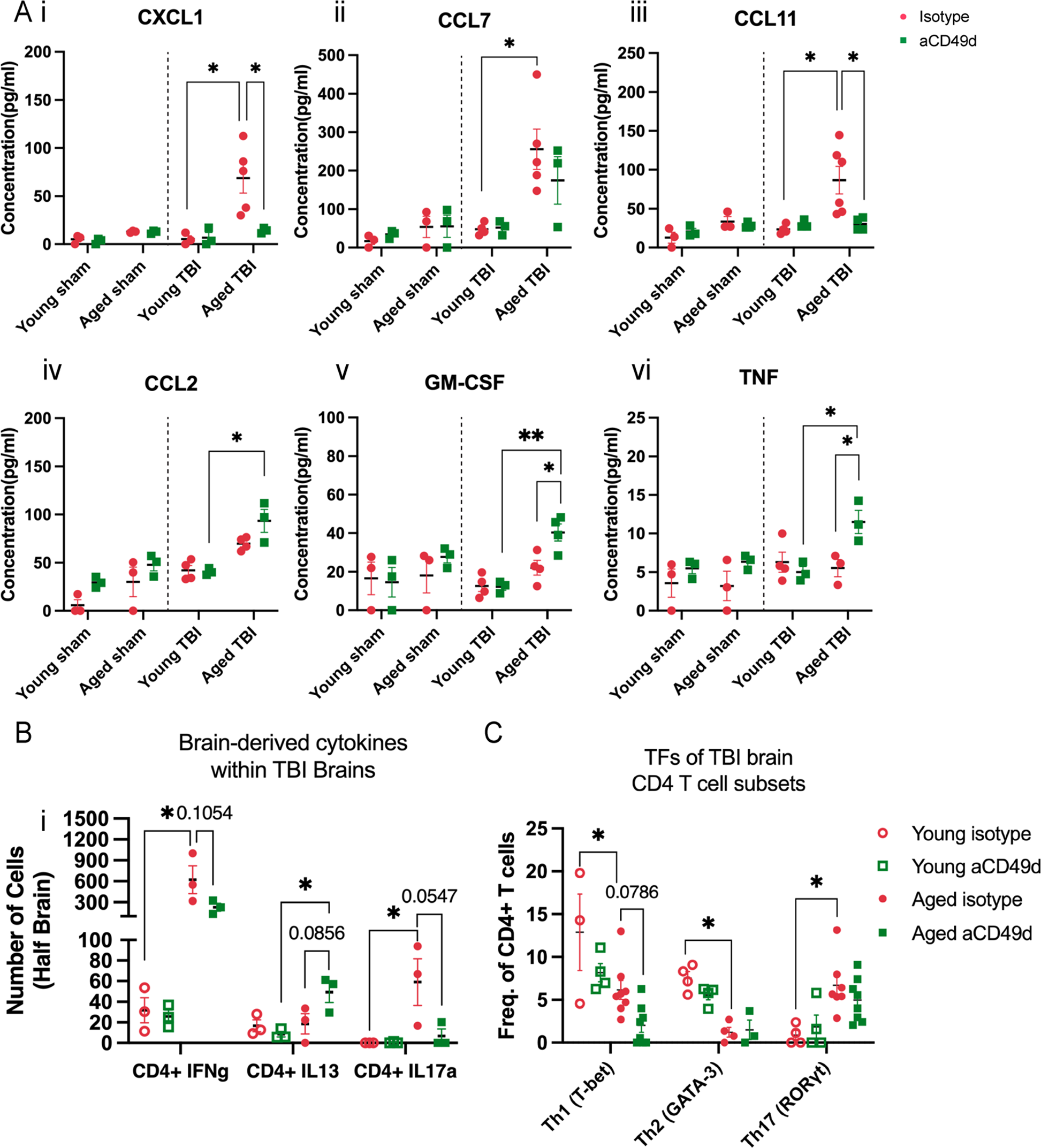
Multiplex cytokine analysis in aged and young mice at 2 months post injury. A. Levels of cytokines in the ipsilateral brain tissue homogenates including i. CXCL1, ii. CCL7, iii. CCL11, iv. CCL2, v. GM-CSF, and vi. TNF-α. B. Cytokine response in brain-derived T cells isolated from aged TBI mice post *ex vivo* stimulation. C. Frequency of Th1, Th2, and Th17 as measured by corresponding transcription factors(TF) T-bet+, GATA-3+, and RORγt+ CD4+ T cells. n =3-8/group, *p < 0.05, **p < 0.01. All data are from one independent experiment. Data are shown as the mean ± SEM, 2-way ANOVA with Tukey’s multiple comparisons test was used for A and the Student’s t test was used for B and C.

### aCD49d Ab Repopulated MHCII high Macrophages within Aged Brains with Chronic TBI

Given the observed GM-CSF elevation in aged TBI mice after aCD49d treatment, we compared the number of macrophages, monocytes, and microglia within aged versus young mouse brains 2 months after TBI. Brain macrophage numbers in aged TBI mice increased dramatically after aCD49d Ab treatment as compared to isotype control (Supp Fig. 5A). Monocytes and microglia numbers remained relatively stable regardless of treatment (Supp Fig. 5B-C). In a separate experiment, we scrutinized the quantity of brain macrophages under different age, injury, and treatment status. Subsequently, we observed an age-associated change in the quantity of macrophages based on MHC status at basal levels: aged sham mice had attenuated MHCII^high^ macrophages (Fig. 4Ai) yet significantly increased MHCII^low^ macrophages (Fig. 4Aii) as compared to their younger counterparts. 2 months post TBI, aged mouse brains had significantly fewer MHCII^high^ but not MHCII^low^ macrophages as compared to young mouse brains (Fig. 4Ai and ii). Repeated aCD49d Ab treatment appeared to upregulate MHCII^high^ macrophages in the aged TBI mouse brains, corresponding to elevated GM-CSF in aged TBI mice with treatment (Fig. 4Ai). To our surprise, aCD49d Ab treatment significantly reduced MHCII^low^ macrophages in the aged sham mouse brains. No other changes were noted after aCD49d Ab treatment in other sham groups. Indeed, recruited monocyte-derived macrophages are capable of differentiating into tissue-specific macrophages and become long-lived CNS boarder-associated macrophages (BAM; 10). Conversely, some BAMs can self-maintain, thereby retaining their yolk sac origin (10). To further decipher their identify, CD45+CD64+ macrophages from aged TBI mouse brains were gated on key surface markers for BAM including CLEC12A, MHCII, P2RX7, MMR, FOLR2, NRP1, CD63 (gating strategy shown in Supp Fig. 6), as described previously (10). As a result, we identified 6 BAM subtypes: MHCII^low^ dural BAM (D-BAM), MHCII^low^ subdural BAM (SD-BAM), MHCII^low^ choroid plexus epiplexus BAM (CP^epi^-BAM), MHCII^low^ choroid plexus BAM (CP-BAM), MHCII^high^ D-BAM, MHCII^high^ CP-BAM. These BAMs reside in different brain compartments as depicted in Fig. 4Bi where MHCII^low^ and MHCII^high^ D-BAMs are found in the dura mater, MHCII^low^ and MHCII^high^ CP-BAMs in the choroid plexus stroma, CP^epi^-BAMs in the apical CP epithelium, and SD-BAms in the subdural. With aging, we noted a reduction in MHCII^high^ D-BAMs (Fig. 4Aiii) and increases in MHCII^low^ D-BAMs, CP^epi^-BAMs, and CP-BAMs (Fig. 4Aiv), consistent with the age-associated changes in MHCII^high^ and MHC^low^ macrophages seen in Fig4Ai and ii. In relation to aCD49d treatment, we observed a nonsignificant increase in MHCII^high^ CP-BAM and MHCII^low^ CP^epi^-BAM in aged mice post TBI as compared to isotype treatment (Fig. 4Aii-iv).

**Figure 4.**
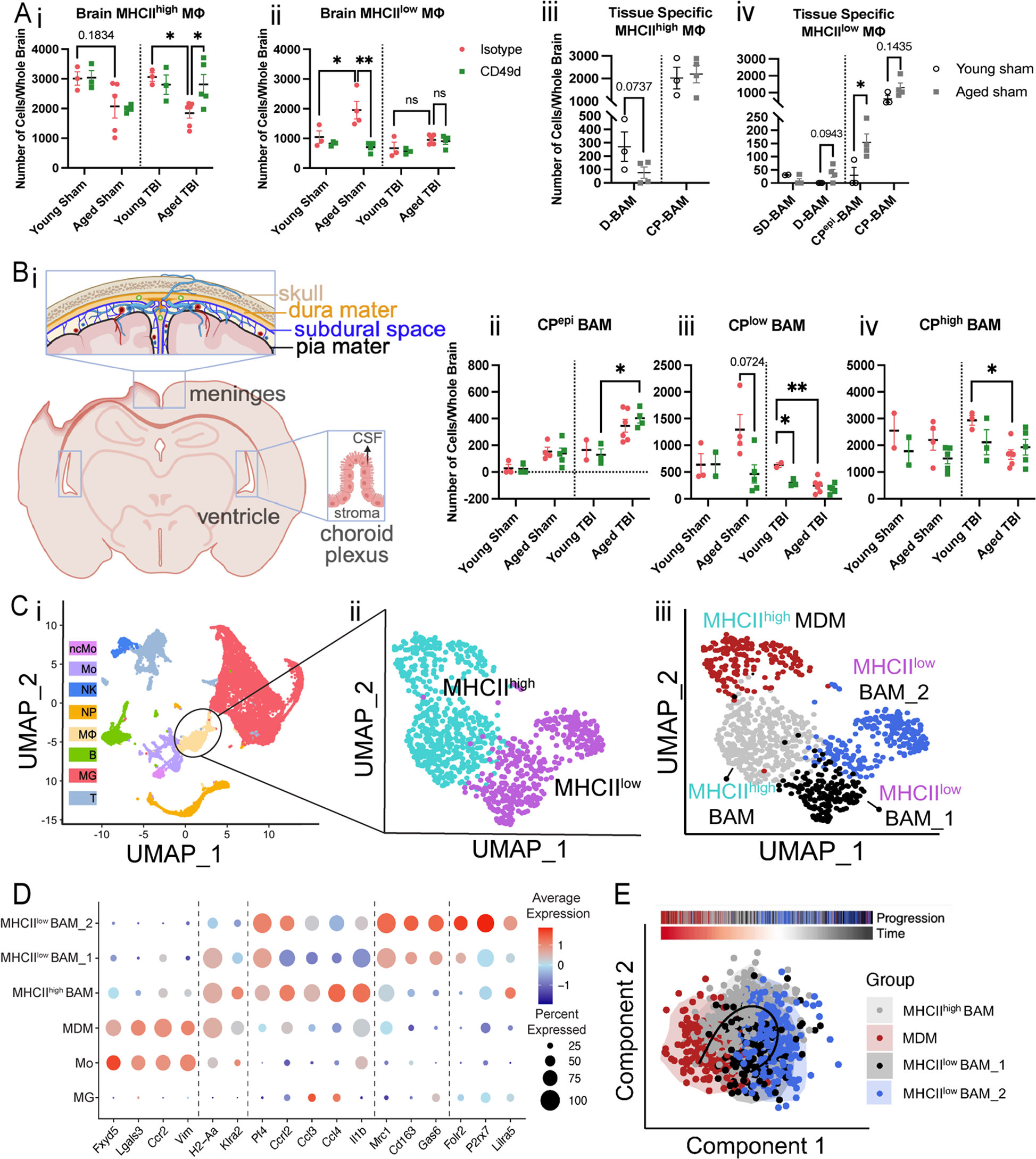
A. Quantifications of i. MHC^high^ and ii. MHC^low^ brain macrophages in both young and aged mouse brains and iii-iv their tissue specificity in sham mice. D-BAM, dural BAM; SD, subdural BAM; CP, choroid plexus BAM. B. i. locations of BAMs in different compartments of the brains. ii-iii. Quantifications of CP BAMs across different sample groups. Data are from one independent experiment and shown as the mean ± SEM, 2-way ANOVA with Tukey’s multiple comparisons test for i and ii, Student’s t-test for iii and iv. n=2-6/group, *p < 0.05, **p < 0.01. C. UMAP plot showing i. CD45+ immune cells isolated from 4 samples (young isotype TBI, young aCD49d TBI, aged isotype TBI, aged aCD49d TBI) for single-cell RNA seq experiment and clustered into 8 cell types by specific markers. ncMO, nonclassical monocytes; Mo, monocytes; NK, natural killer cells; NP, neutrophils; MΦ, macrophages; B, B cells; MG, microglia; T, T cells. ii. UMAP plot showing MHC^high^ and MHC^low^ macrophage cluster and iii. identified subsets within the macrophage cluster. MDM, monocyte-derived macrophages; BAM, CNS boarder-associated macrophages. D. Dot plot showing markers used to identify different macrophage subsets and Itga4 (CD49d) and compare their expression with Mo and MG. E. SCORPIUS trajectory inference on the macrophage cluster.

### aCD49d Ab Treatment Induced Transcriptome Changes in Macrophages in Aged but not Young Brains with Chronic TBI

To investigate the genes, signatures, and pathways involved with aCD49d Ab treatment, we applied single-cell RNA sequencing (scRNA-seq) to brain tissue collected from both young and aged TBI mice treated with aCD49d ab or isotype control (YTBI_Istoype, ATBI_Isotype, YTBI_aCD49d, ATBI_aCD49d). For each group, we pooled two biologic replicates to reduce biologic variance. We excluded sham mouse brains as they had a limited number of infiltrating immune cells. The scRNA-Seq data identified eight distinct cell populations embedded in a Uniform Manifold Approximation and Projection (UMAP) plot (Fig. 4Ci). Focusing on the macrophage population by their MHCII status, we separated the macrophage cluster into MHCII^high^ and MHCII^low^ (Fig. 4Cii). Given that macrophages in the mouse brains exhibited distinct transcriptional states depending on their ontogeny and tissue environment, we further divided macrophages into four subclusters (Fig. 4Ciii and D). The monocyte-derived macrophage (MDM) subcluster was characterized by their monocyte origins and recruitment (*Ccr2*, *Fxyd5*, *Lgals3*, *Vim*), whereas the BAM subcluster universally expresses *Pf4*. Based on recently reported gene markers differentially expressed in MHCII^low^ versus MHCII^high^ BAMs (*Mrc1*, *CD163*, *Gas6* versus *H2-Aa*, *Klra2*), we identified a MHCII^high^ BAM subcluster and two MHCII^low^ BAM subclusters which differed by the expression of signature genes specific for meningeal MHCII^low^ BAMs (*Folr2*), SD-BAMs (*P2rx7*), and CP-BAMs (*Lilra5*) (10). Interestingly, the MHCII^high^ BAM subcluster showed enriched expression of *Ccrl2*, *Ccl3*, *Ccl4*, and *ll1b*, indicating an increased migration ability and inflammation (Fig. 4D).

Significantly upregulated DE genes (adjusted P<0.01, log2(FC)>0.25, present in at least 50% of the cells) in the MDMs within the aCD49d Ab-treated versus isotype-treated aged TBI groups were associated with macrophage activation/polarization including the T cell–interacting activating receptor on myeloid cells 1 (*Tram1*), late endosomal/lysosomal adaptor MAPK and MTOR activator 1 (*Lamtor1*), the receptor for activated protein C kinase 1 (*Rack1*), and eukaryotic elongation factor 1A1 (*eEF1A1*). These genes are associated with cytokine production (Supp Fig. 7Ai and iii). Top DE genes in the MHCII^low^ BAM_2 cluster in aCD49d Ab-treated versus isotype-treated aged TBI groups included *Cd86*, and BAM signature genes *Pla2g7*, *Ninj1*, and *Clec4n* encoding platelet-activating factor acetylhydrolase, ninjurin 1, and C-type lectin domain family 4, respectively, with enriched pathways including negative regulation of activated T cell proliferation (Supp Fig. 7Aii and iii). We did not identify any significant DE genes in the MHCII^low^ BAM_1 cluster between aCD49d Ab-treated and isotype-treated aged TBI groups, and significant DE genes in the MHCII^high^ BAM between aCD49d Ab-treated versus isotype-treated aged TBI groups overlapped with those in the MDM cluster (data not shown). On the other hand, comparing aCD49d Ab treated versus isotype-treated young TBI groups within the four clusters, all DE genes were without significance. When performing trajectory inference, these cells were ordered along a line for inferring developmental chronologies from MDM to either BAM groups (Fig. 4E) with the most predictive genes clustering into modules and depicted in a heatmap (Supp Fig. 8).

### aCD49d Ab Treatment Leads to Fewer Clonally Expanded CD8^+^ T cells in Aged Brains with Chronic TBI

As we previously identified, aCD49d Ab treatment exerts its effect mainly through targeting CD8^+^ T cells. Herein, we integrated single-cell TCRseq with single-cell RNAseq to assess the clonality of T cells and their transcriptomes post-TBI. T cell clusters from the scRNAseq data (Fig. 4Bi) comprised of clonal (C) and nonclonal (NC) T cells (Fig. 5A). We utilized the atlases in Andreatta et al (14) to identify nine T cell subclusters: 1) CD8+ naïve-like (NL) cells with enriched genes including *Cd8a*, *Il7r*, and *Tcf7*, 2) CD8^+^ effector memory (EM) cells high in *Cd8a* and granzymes *Gzmk*, *Gzmb* but low in *Tox*, *Pdcd1*, 3) CD8^+^ early active cells (EA; an intermediate CD8^+^ profile between NL and EM, 4) CD8^+^ terminally exhausted (Tex) cells with high expression of granzymes and inhibitory receptors (*Pdcd1*, *Ctla4*, *Lag3*) and *Tox*, 5) CD8^+^ precursor-exhausted (Tpex) cells enriched in inhibitory receptors but low in granzymes, 6) CD4^+^ NL cells enriched in *Cd4*, *Il7r*,*Ccr7*, and *Tcf7*, 7) CD4^+^ follicular-helper (Tfh) high in *Cd4* and *Tox*, 8) Th1 (*Cd4* and *Fasl*), and 9) Treg cells (*Cd4* and Foxp3) (Fig. 5B and C). Overall, T cell cluster was made of CD8^+^ NL (31.6%) and CD8^+^ EM cells (31.9%) with only few CD8^+^ Tex and Tpex cells (1.6% and 1.9%) (Fig. 5D). As expected, most clonally expanded T cells were CD8^+^ EM cells (72.4%), followed by CD8^+^ NL cells (48.3%) and CD8+ EA cells (44.0%) (Fig. 5D). By groups, T-cells from Aged TBI-Isotype treated mice had a higher proportion of clonal T cells than did the Aged TBI-aCD49d ab treated group (72.5% versus 62.3%; Fig. 5Ei). This was consistent in CD8^+^ EM (93.8% versus 82.9%; Fig. 5Eii) and CD8^+^ NL cells (70.0% versus 59.1%; Fig. 5Eiii). On the contrary, no such difference was seen in aCD49d treated young mouse brains (Fig. 5Ei-iii). To delineate the relative abundance of TCR clones in their individual repertoire, we applied scRepertoire R package (15) to quantify the proportion of clones by their size. As a result, 14.0% of TCR clonotypes in the Aged TBI-Isotype treated samples belonged to hyperexpanded clones (clones taking up more than 1/10 of clonal space) which were not found in the Aged TBI-aCD49d treated samples. This indicates that aCD49d Ab treatment might attenuate excessive clonal expansion.

**Figure 5.**
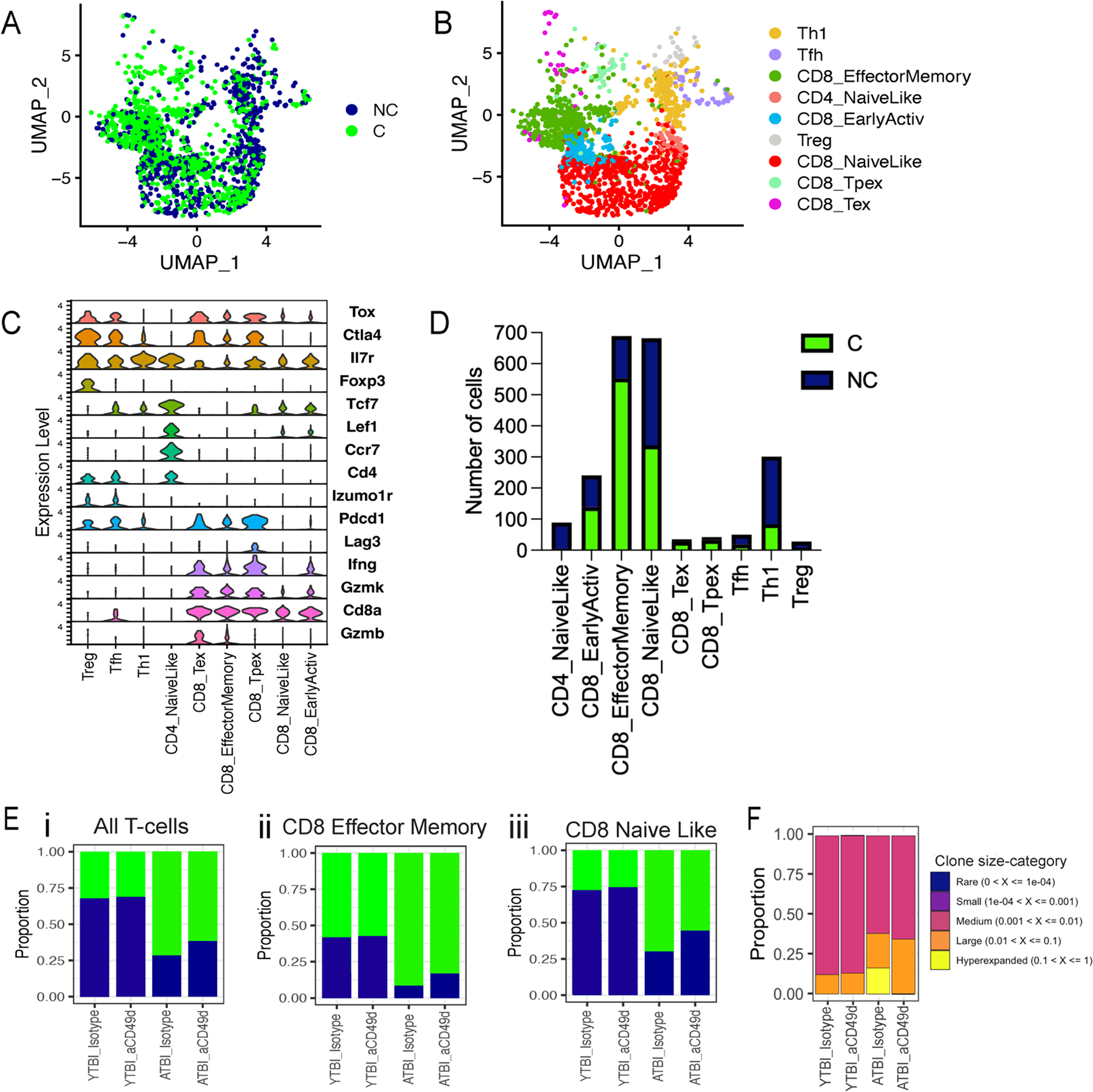
Clonally expanded CD8^+^ T cells patrolled aged mouse brains post-TBI and aCD49d Ab treatment induced transcriptional changes largely in CD8^+^ T Cells. A. single-cell TCR analysis overlaid on UMAP projections showing distribution of T cell clonality. C: clonally expanded. NC: nonclonally expanded. B. UMAP showing distribution of T cell subtypes based on T cell atlases ^4^. C. Dot plot showing the markers to identify different T cell subtypes. D. Bar graph demonstrating number of clonality across T cell subtypes. E. Bar graph demonstrating proportion of i. all T-cell, ii. CD8 effector memory T-cell, and iii. CD8 naïve like T-cell clonality across samples. F. The proportion of clones belong to each clone size category. ATBI: aged TBI mouse brains. YTBI: young TBI mouse brains.

### aCD49d Ab Treatment Reduces CD8+ T cells Exhibiting Higher Effector Status in Aged Mouse Brains with Chronic TBI

The CD8+ T cells (*Cd8a > 1.5*) were further annotated based on various functional markers, followed by an unsupervised clustering (Supp Fig. 9A). Most CD8+ T cells originated from aged TBI mouse brains (i.e., ATBI_isotype and ATBI_aCD49d) with treatment-dependent cluster 6 uniquely expressed in ATBI_aCD49d and cluster 0 in ATBI_isotype. Upon comparing the expression of genes associated with effector, exhaustion, and memory along with clonality (Supp Fig. 9B and C), we found that aged mouse brains had a substantial number of dysfunctionally activated CD8+ T cells, herein defined as those that highly co-express genes associated with exhaustion (*Pdcd1*, *Entpd1*, *Ctla4*, *Havcr2*) and effector genes (*Gzmk*, *Gzmb*, *Prf1*, *Tnf*, *Ifng*). We also found two clusters of treatment-dependent functionally activated CD8+ T cells that are high in effector genes only, and two clusters of age-dependent stem-like memory T cells (*Tcf7*, *Slamf6*, *Lef1*, *Sell*) (Fig. 6A). Nonclonal T cells high in *Ccr7* were defined as naïve-like T cells (Supp Fig. 9B, Fig. 6A). *Isg15* marked a cluster of IFN dominated CD8+ T cells. Lastly, we found a small cluster with CD8+ T cells in S phase only (Fig. 6A). Thereafter, aCD49d Ab decreased functionally activated CD8+ T cells proportionally compared to isotype treatment (6.6% versus 29.8%; Fig. 6Ai and Aii). Dysfunctionally activated and functionally activated CD8+ T cells are transcriptionally distinct as indicated by significant DE genes associated with exhaustion (*Ctla4*, *Pdcd1*), migration (*Ccl4*, *S100a11*), and tissue-residency (*Cxcr6*) (Fig. 6Bi). Contrastingly, functionally activated CD8+ T cells differentially upregulated effector gene *Ly6c2*. Flow cytometry further validated that aCD49d Ab treatment significantly reduced activation as demonstrated by PD-1 production in CD8^+^ T cell as compared to isotype treatment (Supp Fig. 10Ai and ii). Meanwhile, aCD49d Ab treatment showed a nonsignificant reduction in CD8^+^ T cell activation marked by CD69 expression (Supp Fig. 10Bi and ii). Furthermore, CD49d^high^ CD8+ T cells expressed significantly increased expression of granzyme B than did CD49d^low^ CD8+ T cells within TBI mouse brains (Fig. 6Ci and ii). Similarly, CD49d^high^ CD8+ T cells expressed significantly increased expression of Ly6C than did CD49d^low^ CD8+ T cells within TBI mouse brains (Fig. 6Ci and iii). aCD49d Ab treatment significantly %CD49d^high^ CD8+ T cells and %Granzyme B^+^ CD8+ T cells in aged mice after 2 months post TBI (Fig. 6D i and ii). aCD49d Ab treatment also led a nonsignificant reduction of %Ly6C^+^ CD8+ T cells in aged mice after 2 months post TBI (Fig. 6D iii).

**Figure 6.**
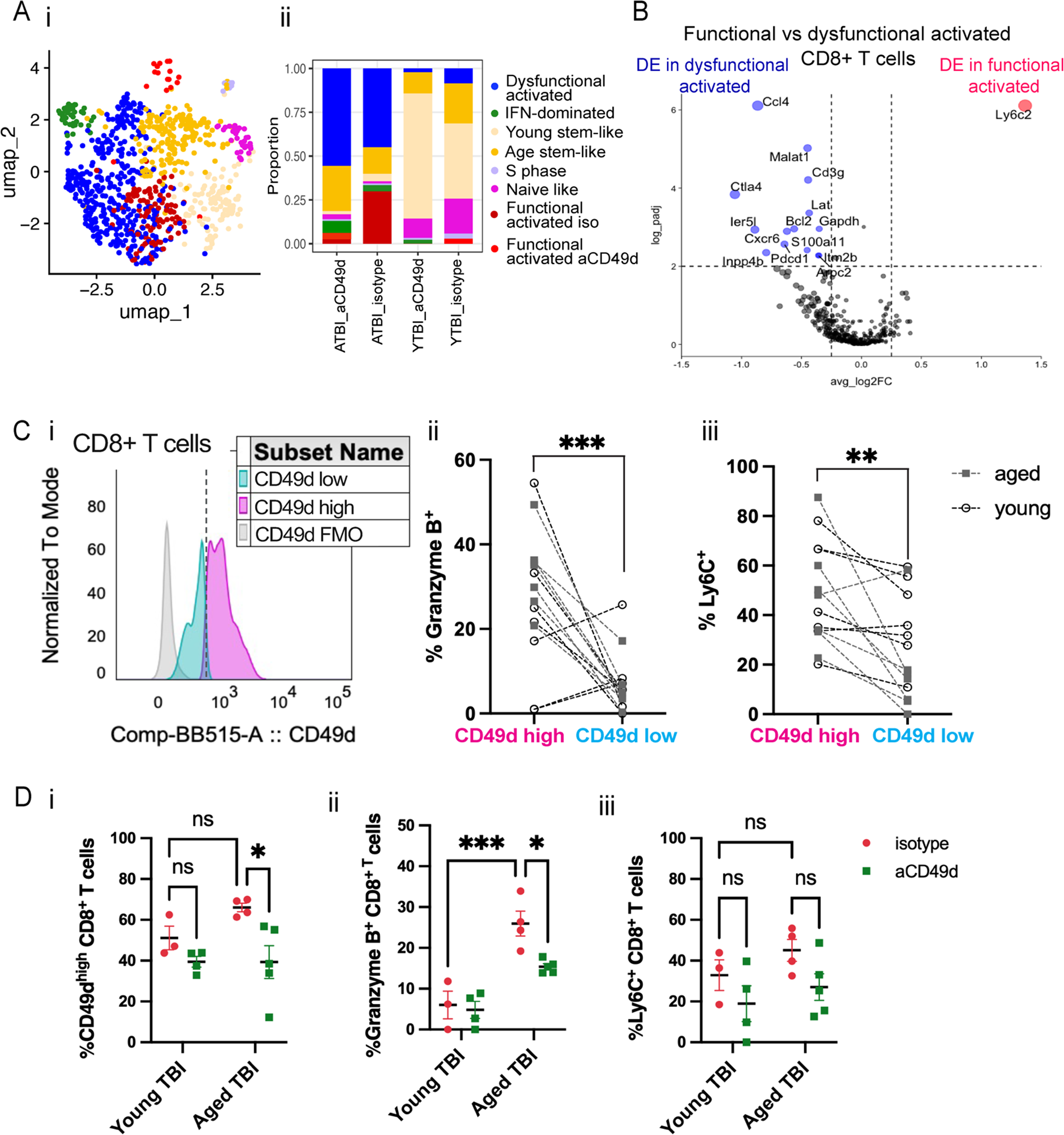
aCD49d Ab treatment reduces CD8+ T cells that exhibit higher effector status in aged mouse brains after TBI. A. i. UMAP showing distribution of CD8+ T cell subtypes based on various T cell differentiation states. ii. Bar graph demonstrating proportion of different T cell differentiation states across samples. B. Volcano plot depicting DEGs in functional vs dysfunctional activated CD8+ T cells within the brains of aged, injured mice two months post TBI. ATBI: aged TBI mouse brains. YTBI: young TBI mouse brains. C. i. Histogram showing cutoff for CD49d high vs CD49d low in CD8+ T cells. ii. Comparison of % granzyme B high in CD49d high vs CD49d low CD8+ T cells. iii. Comparison of % Ly6C high in CD49d high vs CD49d low CD8+ T cells. D. i. % CD49d high in CD8+T cells across TBI samples. ii. % granzyme B high in CD8+ T cells across TBI samples. iii. % Ly6C high in CD8+ T cells across TBI samples. Data are from one independent experiment and shown as the mean ± SEM. 2-tailed paired student’s t test, n=12 for C. 2-way ANOVA with Tukey’s multiple comparisons test for D, n=3-5/group, *p < 0.05, **p < 0.01, ***p < 0.001.

## Discussion

Immediately upon TBI, parenchymal microglia converge upon the injury and initiate a cascade of immune events in conjunction with the surrounding astrocytes (16, 17). This immune cascade releases cytokines and chemokines to recruit peripheral innate immune cells to lesion areas, serving as the first line of defense to resolve injury and remove cellular debris. Indeed, numerous cytokines and chemokines are produced by systemic leukocytes, glial cells, and even neurons, all acting in concert as critical regulators of the injury process (18). Though traditionally viewed as pro- or anti-inflammatory in action, these regulators are now classified based on their receptor interactions with a net effect aiming to limit the spread of the injury and restore homeostatic balance (19). Unfavorable TBI outcomes have been linked to five major mediators including IL1, IL6, TNF, TGFβ, and IL10 as well as neutrophil chemokines CXCL1/2 (19–21). Poorer prognosis for older TBI mice is partially attributable to markedly more IL1, IL6, IL22 (part of IL10 family), TNFα and CXCL1/2 in both the periphery and with the injured brain as compared to their younger counterparts after acute TBI (Fig. 2, Supp Fig. 4). Furthermore, at 2 months post-TBI aged mice express elevated chemoattractants including CCL7 and CCL11 in their brains, with CXCL1 remaining high (Fig. 3). Chronically high neutrophil chemokines lead to vigorous neutrophil recruitment. Along with other infiltrating leukocytes, these over-accumulating peripheral innate immune cells release toxic molecules including pro-inflammatory cytokines, free radicals, and proteases within the injured brain, causing toxicity to neurons and promoting secondary tissue damage (19, 22, 23). By deciphering the age-specific differential cytokine and chemokine response after TBI, our study revealed that aCD49d Ab treatment could markedly attenuate these age-associated, heightened, and toxic cytokine and chemokine responses post TBI. This, in turn, improved outcomes in aged mice (Fig. 2 and 3, Supp Fig. 4).

Our findings are consistent with recently published data showing a T helper 1/T helper 17 (Th1/Th17) polarization in the blood and with the injured brain up to 3 weeks after TBI, preceding CD8^+^ T cell infiltration during the chronic phase (24, 25). Moreover, compared to young mice, aged mice further upregulated Th1/17 immunity as indicated by elevated IL12(p70) and IL17A levels within the first week post-TBI in both the blood and within the injured brain (Fig.2 and Supp Fig. 4A; 26). Th1 and Th17 responses exaggerate inflammation (27). The Th17 response can further enhance the cytotoxicity of CD8^+^ T cells once infiltrated into the injured brain thereby accelerating neuroinflammation and, ultimately, lead to demyelination (25, 28, 29). Meanwhile, the Th2 response is neuroprotective and is reduced after TBI (25, 27). Our data show that aCD49d Ab treatment significantly mitigated the increased Th1/Th17 polarization in aged mice during acute TBI and augmented a neuroprotective Th2 polarization with upregulation of IL-4 and IL-13 in the late phase of TBI (Fig.2, Fig. 3Bii, and Supp Fig. 4A; 8). Additionally, we cannot exclude the possibility of the Th2 response being derived from the group 2 innate lymphoid cells (ILC2s), as they can also produce IL-4 and IL-13. Furthermore, ILC2s are also known to produce GM-CSF (28, 29). In the meantime, aCD49d Ab treatment markedly blocked CD8^+^ T infiltration as indicated in the snapshot of CD8^+^ T infiltration measured in the *in vivo* CD45^+^ labeling and the total number of CD8^+^ T in the brain parenchyma post TBI in aged mice (Fig. 1F and H). The augmented Th2 response by aCD49d Ab during the chronic phase of TBI is likely due to attenuated cytotoxicity of CD8^+^ T cells as similar results were reported in genetic deficiency or pharmacological depletion of CD8^+^ T cells (25). It is important to note that the depletion of CD4^+^ T cells does not improve neurological outcomes post TBI (25). Hence, the age-specific therapeutic effects observed in repeated aCD49d Ab treatment is largely attributed to the blockade of CD8+ T cell infiltration and suppression of the age-associated, heightened, and toxic cytokine and chemokine response.

Normal aging results in decreased function of T cells, including reduced proliferation, reduced survival potential, short-lived effector T cells, and diminished long-term T-cell memory (30). Furthermore, previously published data have shown that aging results in increased numbers of CD8+ T cells within the brains of both mice and humans (31). Recent studies have shown that the age-associated CD8+ T cell dysfunction in mouse brains is likely related to T cell senescence, which is driven by elevated cytokines rather than specific antigens and displays a proliferation defect (32, 33). Another study reported that CD8+ T cells expressing the programmed cell death protein 1 (PD-1), indicative of CD8+ T cell dysfunction, accumulate in the aged mouse brain (34). Our findings revealed that compared to young mice brains, aged mouse brains have two pools of CD8+ T cells: those which constantly infiltrate from the blood and those which have acquired residency but are dysfunctional. Our findings posit that aCD49d Ab exert its therapeutic effects through targeting the pathologic CD8+ T cell infiltration (Fig. 6Di), in turn leading to a reduction of functionally activated CD8+ T cells within the aged mouse brains after TBI (Fig. 6A and B). Repeated aCD49d Ab treatment led to a profound reduction of PD-1^+^ and Granzyme B^+^ CD8+ T cells within the brains of aged mice as compared to isotype control-treated mice (Supp Fig. 10 and Fig. 6Dii), which resulted in a less clonal expansion throughout chronic TBI (Fig. 5E and F). In agreement with prior work in a cancer model, we found that CD49d^high^ CD8+ T cells exhibit greater cytotoxic potential than CD49d^low^ CD8+ T cells within TBI mouse brains as indicated by Granzyme B and Ly6C expression (Fig. 6Cii and iii). In TBI, CD8+ T cells have been found to exacerbate the index injury and cytotoxic CD8+ T cells have been documented to cause direct cytotoxic effects on neurons (35–38). Collectively, aCD49d Ab blockade of the α4 integrin may represent a novel treatment strategy in age-associated traumatic brain injury.

Monocytes represent a major element of this cellular infiltration. They enter into the injured brain, giving rise to macrophages and contributing to scar formation, tissue resolution, and acute neuronal death (39–41). Akin to current literature, our study showed a portion of the overall border-associated macrophages (BAM) population being replenished by monocyte-derived macrophages (MDM). In contrast, some BAMs can self-repopulate depending on their ontogeny (10–12). aCD49d Ab treatment significantly upregulated whole-brain GM-CSF (Fig. 3Av) and increased the overall macrophage population in the aged mouse brain 2 months post TBI (Supp Fig. 5A). Meanwhile, aCD49d Ab treatment had major effects on MHCII^high^ macrophages within the aged brains but little effect on MHCII^low^ macrophages at 2 months post TBI (Fig. 4Ai and ii). Thus, it appears that aCD49d Ab treatment facilitates the turnover of MHCII^high^ BAM by promoting the conversion of MDMs into BAMs. Based on our findings, we surmise that MHCII^high^ MDM-converted BAMs might further become MHCII^low^ BAMs (Fig. 4E). This potentially explains the decreased MHCII^high^ BAMs and increased MHCII^low^ BAMs that we observed in aged sham mice as compared to young sham mice: MDMs are uniformly MHCII^high^ but might lose MHCII expression when they acquire a BAM profile. Indeed, MHCII expression on macrophages is regulated by environmental stimuli such as the gut microbiome for gut macrophages and their ontogeny (10–12). Given that both aging and TBI contribute to the turnover and function of BAMs, future studies are needed to elucidate the effects of aCD49d Ab on distinct subsets of BAMs.

Transcriptomically, with aCD49d Ab treatment, MDM cluster upregulated essential genes, including Lamtor1 and *eEF1A1,* that have been found to drive the macrophage activation to the tissue repair phenotype, also known as M2-like macrophages (Supp Fig. 7Ai; 42, 43). Furthermore, an augmented Th2 response, indicated by elevated IL-4 and IL-13, polarizes macrophages towards the tissue repair phenotype (44). It is worth noting that CCL2 was slightly increased with repeated aCD49d Ab treatment in aged mouse brains at 2 months post TBI (Fig. 3Aiv). This implies that the recruited monocytes are quickly converted to anti-inflammatory macrophages at the site of injury and participate in the repair process (45, 46). Polarization of a subset of macrophages to a M2-like anti-tumor phenotype has been observed in glioblastoma and lung cancer (47, 48). Our study was limited due to the nature of single-cell RNAseq. Further experiments such as bulk RNAseq, which has a higher sequencing depth, and quantitative real-time PCR will be helpful to validate our DE findings. Additionally, the meninges were not included in our analysis, where a subpopulation of BAMs reside. This explains the limited number of dural and subdural BAMs, as displayed in Fig.4 Aiii and iv. Given that CCI is an invasive model of TBI whereby the meninges are broken down, it is also possible that BAMs extravasated into the brain parenchyma allowing for their detection. Our findings pinpointed that aging, TBI, and MHCII class status collectively impact the complexity and heterogeneity of brain macrophages, including BAMs. Further characterization incorporating these factors is critical and can complement our findings as well as deepen the understanding of brain macrophages, a highly heterogeneous population of cells that are orchestrated to perform various specialized functions.

The US population is aging at an unprecedented rate with the projected number of Americans aged 65 years and older to be greater than 50% by 2050 (49). TBI in older adults is a growing public health concern that requires a focused, age-specific, approach. As such, our findings have high clinical relevance for treating TBI in older patients. aCD49d Ab (Natalizumab) has undergone phase 1, 2, and 3 clinical trials and is currently FDA approved for use in acute stroke, relapsing forms of multiple sclerosis, and Crohn’s disease-induced inflammation (50–53). Unlike reports stating no effect was seen in the chronic phase of stroke, aCD49d Ab improved functional outcomes during the chronic phase (2 months) of TBI in our model (50, 54). Compared to antiCD8 Ab therapy, which depletes CD8^+^ T cells entirely, aCD49d Ab does not lead to long-term immunosuppression and could be administered intravenously as soon as a diagnosis of TBI is made in the geriatric trauma patient. Additionally, ACD49d Ab therapy has the potential to beneficial in other age-associated neurodegenerative diseases associated with CD8^+^ T cell infiltration, such as Alzheimer’s disease and Parkinson’s disease (55, 56).

We have previously published that blocking the infiltration of CD8+ T cells with aCD49d Ab improved survival, neurocognitive outcomes, and motor function in aged mice after TBI (8). The current study shows that the mechanisms underlying this protection include a significant attenuation of CD8^+^ T cell cytotoxicity, an age-associated, heightened, and toxic cytokine and chemokine response, along with an augmented neuroprotective Th2 response. In summary, this study comprehensively examined the putative mechanisms leading to worse TBI outcomes aged subjects. It adds substantially to our understanding of aging as a significant risk factor for poor outcomes after brain trauma. In light of these findings, future clinical trials should consider aCD49d Ab as a novel, age-specific, treatment for TBI and other neurodegenerative disease processes.

## Methods

### Mice

C57BL/6 mice were used in all experiments with aged mice from National Institute on Aging (NIA) aged rodent colonies (Gaithersburg, MD) and young mice from Jackson Laboratory (Bar Harbor, ME). All mice were housed in a barrier facility at the Northwestern University Center for Comparative Medicine (Chicago, IL). Young adult mice underwent TBI versus sham injury at 13-15 weeks of age (n = 25), and aged mice underwent TBI versus sham injury at 80-82 weeks of age (n = 31). Animals were housed in ventilated cages under standard conditions at a temperature of 24°C and a 12-hour light/dark cycle with ad libitum access to standard food and water.

### Sex as a biological variable

Our study examined male mice because female animals, due to the estrous cycle, exhibit in differential TBI outcomes.

### Controlled cortical impact

TBI or sham injury was induced via controlled cortical impact (CCI), as previously published by our laboratory (7,8,57). Animals were anesthetized with 50 mg/kg Ketaset Ketamine (Fort Dodge, IA) and 2.5 mg/kg xylazine (AnaSed; Shenandoah, IA) via intraperitoneal injection. All mice underwent a medial 1-cm longitudinal incision to expose the skull. TBI mice underwent a 5-mm craniectomy located 2 mm to the left of the sagittal suture and 2 mm rostral to the coronal suture over the M1/S1 cortex. The dura mater was intact. TBI mice then received a 2-mm depth injury delivered by a 3-mm impacting tip at 2.5 m/s with a 0.1-second dwell time. The incisions were closed using VetBond 3M (Santa Cruz Animal Health, Dallas, TX). Sham animals underwent anesthesia and scalp incision only. Postoperative analgesia (0.1 mg/kg buprenorphine SR) was administered to all mice.

### Anti-CD49d antibody treatment

Mice were treated with 300 µg anti-CD49d (BE0071, clone: PS/2; Bio X Cell) or isotype control (BE0090, clone: LTF-2; Bio X Cell) in *InVivo*Pure Dilution Buffer (pH 6.5 for anti-CD49d Ab or 7.0 for isotype control) 2 h after CCI or sham surgery, followed by injection every second week until termination of the experiments as we have previously published (8).

### 5-ethynyl-2’ -deoxyuridine (EdU) administration

Mice were treated with EdU (5 mg/ml; invitrogen) in PBS (pH 7.4) daily via intraperitoneal injection at a dose of 50 mg/kg body weight for 3 days before saline perfusion. 50 mg/kg EdU is the saturating dose and is adequate for labeling proliferating cells at near saturation levels in the adult mouse brains (58). EdU was detected using a Click-iT EdU Alexa Fluor 647 Flow Cytometry Assay Kit (C10635; Invitrogen) following the manufacturer’s instructions.

### Intravascular labeling

For intravascular labeling experiments, 200uL total volume of 3ug or 15uL of BUV661 CD45 (612975, BD Biosciences) in 285uL PBS was injected via retroorbital injection 2 hours before euthanasia.

### Behavioral testing

All tests were performed in the Northwestern University Behavioral Phenotyping Core during the light phase cycle. Experiments and analyses were performed in a blinded fashion with regard to surgery status and treatment.

### Y maze assay of spontaneous alternation

Y maze testing was conducted 30 days post-TBI to measure the spatial working memory (59). The mouse was placed in a Y-shaped maze with three arms of identical dimensions at a 120° angle from each other. The mouse was introduced at the base of the arm forming the stem of the Y with its nose towards the center of the maze and allowed to explore the arms freely. The mouse was observed for five minutes, and its movement was recorded. The order of arm entries was analyzed for spontaneous alternation (SA). The box was cleaned with 70% ethanol in between trials to minimize olfactory cues. The video was captured and analyzed by LimeLight 4 software (Actimetrics, Wilmette, IL).

### Rotarod

Rotarod was conducted 45-50 days post-TBI to measure the motor coordination (60). The mice were placed onto a five-lane rotarod device with the latency to fall (s) being recorded. Five mice ran together per trial (4 trials per day) and underwent three-day habituation trials at a constant speed of 12RPM for a maximum of 5 min. On day 4-5 acceleration trials, rotarod accelerated from 4-40 RPM over 5 min. The trials ended when the mouse fell from the rod. The equipment was cleaned with 70% ethanol in between trails to eliminate olfactory cues. Individual scores from four trials were averaged and evaluated between different injury and treatment groups.

### Tissue harvesting and blood collection

For the experiment in Supplement Figure 3, blood sampling via submandibular blood collection was performed using GoldenRod animal lancet (Medpoint Inc). For all rest experiments, depending on the experimental approach, mice were euthanized at either 3 days or 2 months post injury via carbon dioxide inhalation consistent with the AVMA guidelines. Blood was collected from euthanized mice via cardiac puncture using a 25-gauge syringe. Collected blood was transferred into an EDTA tube and then centrifuged for 15 min at 3000 rcf for plasma collection. Plasma samples were stored at −80°C until analysis. The cadavers were then perfused through the aorta with 20 mL of 4°C 1x Hank’s balanced salt solution (HBSS). Brains were excised and placed in ice-cold HBSS until processing.

### Preparation of brain homogenates and brain cells

Brains were separated into ipsilateral and contralateral hemispheres. 500 µL of cell lysis buffer (EPX-99999-000; Invitrogen) was added to each hemisphere in M-tubes followed by homogenization with a MACs dissociator (Miltenyi Biotec, Inc) according to the manufacturer’s instructions. The samples were centrifuged at 16,000 g for 10 min at 4°C. The supernatants were transferred to new microcentrifuge tubes and stored at −80°C until analysis. Additionally, brain cells were isolated as we have previously described (7, 8). Cell debris was removed using the Debris Removal Solution (130-109-398; Miltenyi Biotec) and a single cell suspension was made. For sequencing experiments, cells were washed and sorted. CD45^+^ cells were sorted on a BD FACSAria cell sorter (BD Biosciences, San Jose, CA) 2 months post-TBI or sham surgery using our previously published gating strategy (7).

### Multiplex cytokine assay

Cytokine profiling of plasma and brain tissue homogenates was performed using the Th1/Th2 Cytokine & Chemokine 20-Plex Mouse ProcartaPlex kit (EPX200-26090-901; Invitrogen) and measured Th1/Th2: GM-CSF, IFNγ, IL-1β, IL-2, IL-4, IL-5, IL-6, IL-12p70, IL-13, IL-18, TNFα and chemokines: eotaxin (CCL11), GRO alpha (CXCL1), IP-10 (CXCL10), MCP-1 (CCL2), MCP-3 (CCL7), MIP-1α (CCL3), MIP-1β (CCL4), MIP-2, RANTES (CCL5). Samples were assayed according to the manufacturer’s instructions and then read on a Luminex 200 instrument (Luminex Corporate, Augtin, TX). Analysis was performed using Thermo Fisher’s Procarta Plexanalyst application to determine the concentration (pg/ml) of each cytokine/chemokine based on the standard curve.

### Flow cytometry

Cell surface staining was performed using the following antibodies: Fixable Viability Dye eFluor 506 (65-0866-14; Invitrogen), CD45 BUV395 (564279, clone: 30-F11; BD Biosciences), CD11b BV421, CD3 FITC(11-0032-82, clone: 17A2; Invitrogen), CD8 PerCP-Cy5.5(1299159, clone: 53-6.7; BD Biosciences), CD4 AF700(557956, clone: RM4-5; BD Biosciences), CD64 BV786 (clone: X54-5/7.1, 741024; BD Biosciences), I-A/I-E PE-Cy7 (clone: M5/114.15.2, 107629; BioLegend), CD11b-APC-Cy7(557657, clone: M1/70; BD Biosciences), CD206(MMR) BB515 (clone: C068C2, 141703; BioLegend), CD11b APC-Cy7 (clone: M1/70, 557657; BD Biosciences), CD304 (Neuropilin-1/NRP1) BV421 (clone: 3E12, 145209; BioLegend), FOLR2 PE (clone:10/FR2, 153303; BioLegend), CD63 PE-CF594 (clone: NVG-2, 143913; BioLegend), P2X7R PE-Cy7 (clone: 1F11, 148707; BioLegend), I-A/I-E AF700 (clone: M5/114.15.2, 107622; BioLegend), CD11c BUV737 (clone: HL3, 612796; BD Biosciences), CD371(CLEC12A) APC (clone: 5D3/CLEC12A, 143405; BioLegend), CD279 (PD-1) eFlour 450 (clone: J43, 48-9985-80; Invitrogen). For intracellular staining, Foxp3/Transcription Factor Staining Buffer Set (00-5523099; eBioscience) was used following the provider guidelines. The following antibodies for intracellular staining were used: Nur77 PE (clone: 12.14, 12-5965-80; Invitrogen), TOX PE (clone: TXRX10, 12-6502-80; Invitrogen). For cytokine assessment of Th cell subtypes, isolated brain cells were stimulated with 50ng/ml phorbol 12-myristate 13-acetate (PMA), 1μg/ml ionomycin, and 10 μg/ml brefeldin A (MilliporeSignma) for 4 h at 37°C. Following the incubation, cells were stained for surface markers including Live/Dead APC-Cy7, CD4 Pacific Blue, CD8 APC, for 30 min at 4°C. Subsequently, brain cells were fixed and permeabilized with Fixation/Permeabilization buffer kit (88-8824-00; eBioscience) for 30 min at room temperature. Cells were then incubated with IL-17A FITC (clone: eBio17B7, 11-7177-81; Invitrogen), IL-13 PE (clone: eBio13A, 12-7133-41; Invitrogen), and IFN-γ PerCP-Cy5.5 (clone: XMG1.2, 560660; BD Biosciences) for 30 min at room temperature. For all experiments, 123count eBeads Counting Beads were added for quantification of absolute cell numbers (01-1234-42; Invitrogen). Stained cells were then analyzed on a BD FACSymphony flow cytometer (BD Biosciences), and analysis was performed using flowJo software (version 10.0).

### scRNA and TCRseq library preparation and sequencing

The concentration and viability (>85%) were confirmed using a K2 Cellometer (Nexcelom Bioscience LLC, Lawrence, MA) with the AO/PI reagent, and ∼5,000 cells were loaded on 10x Genomics Chip K using the 10x Genomics Chromium Next GEM Single Cell 5’ v2 with immune profiling kit and Controlled (10x Genomics, Pleasanton, CA). Libraries were prepared according to 10x Genomics protocols. Libraries were sequenced on an Illumina NovaSeq 6000 instrument (Illumina, San Diego, CA). Bases were called using the Illumina RTA3 method. RNA reads were aligned to the mm10 mouse reference genome and gene expression matrices were generated using Cell Ranger (version 7.2.0). TCR reads were also aligned to the mm10 mouse reference genome and clonotype/contig matrices were generated using Cell Ranger.

### Processing of scRNAseq and TCRseq data

Gene expression matrices were loaded and read into R using the functions Read10x and CreateSeuratObject in the Seurat R package (version 5.0.1; 61) from the Satija Lab. Background contamination was corrected using R package SoupX (version 1.6.2; 62). Further quality control was performed as we have previously described (7). TCR clonotype/contig matrices were also filtered for empty droplets using Cell Ranger. Only TCR sequences associated with T cells were retained in downstream analysis. Corrected and filtered gene expression matrices were SCTransformed and regressed out based on number of reads and features as well as mitochondrial mapping percentage on a per sample basis using Seurat package. Following that, four groups (YTBI_isotype, YTBI_aCD49d, ATBI_isotype, and ATBI_aCD49d) were integrated as recommended for cell type identification using Seurat. Annotation of identified clusters was performed using canonical markers and assisted by SingleR package (version 2.2.0; 63). The R packages ggplot2 (version 3.4.4) and Seurat were used to plot all plots.

### Downstream bioinformatic analysis

Seurat FindMarkers was used to identify differentially expressed genes (DEGs). Model-based analysis of single-cell transcriptomics (MAST) was chosen to test significance. P values were adjusted for multiple comparisons using the Benjamini-Hochberg procedure. Genes with an adjusted p value less than 0.01 and average log-fold change magnitude greater than 1 were considered significantly differentially expressed. Downstream gene ontology (GO) enrichment analysis was performed using GOrilla with all genes being expressed in the samples as a background. The false discovery rate (FDR) was calculated to correct for multiple tests. SCORPIUS (version 1.0.9) trajectory analysis was performed on macrophage cluster using the proposed workflow with the Spearman correlation being selected (64). R packages ProjecTILs (version 3.3.0) was used to classify T-cell states and embed our results into reference single-cell maps (14). Identified T-cell states were overlaid with T cell clonality from TCRseq for further differential expression analysis of clonally expanded or individual T cells. scRepertoire R package (version 2.0.0) was used to examine the TCR clonal space.

### Statistical analyses

For behavioral tests, a two-way ANOVA with Turkey multiple comparisons test was performed to determine the statistical significance between aged sham with isotype, aged sham with aCD49d Ab, aged TBI with isotype, and aged TBI with aCD49d Ab for the following variables: injury and treatment. The same statistical tests were performed in young mice. For cytokine measurements, infiltrating/proliferating immune cell phenotyping, and brain macrophage quantification, a two-way ANOVA with Turkey multiple comparisons test was performed to determine the statistical significance between young TBI with isotype, young TBI with aCD49d Ab, aged TBI with isotype, and aged TBI with aCD49d Ab for the following variables: age and treatment. For quantification of infiltrating/proliferating/total T cells, Th1/Th2/Th17 T cell response, PD-1/Nur77-expressing CD8+ T cells, and tissue-specific brain macrophages, a student’s t-test was performed. P was set at P < 0.05, using GraphPad Prism (version 6.01; GraphPad Software Inc USA). Outlier boundaries are 1.5 standard deviations away from the mean of the values.

### Study approval

All procedures were approved by the Northwestern University Institutional Animal Care and Use Committee. All animal experiments were performed and reported in accordance with the ARRIVE guidelines.

## Acknowledgments

The work by the authors is supported through the National Institute of Neurological Disorders and Stroke (NINDS), NIH (R01 NS127865), National Institute of Aging (NIA), NIH (T32 AG020506), and Diana Jacobs Kalman/American Federation for Aging Research (AFAR) Scholarships for Research in the Biology of Aging. We thank the Mechanisms of Aging & Dementia Training Program and the Center for Human Immunobiology at Northwestern University for giving scientific advice. We thank University Robert H. Lurie Comprehensive Cancer Center of Northwestern University in Chicago, IL, for the use of the Flow Cytometry Core Facility. The Lurie Cancer Center is supported in part by an NCI Cancer Center Support Grant #P30 CA060553., the NUSeq Core. This work was supported by the Northwestern University NUSeq Core Facility. We would also like to acknowledge NIH Grant 1S10OD025120 for the 10x Chromium housed in the NUSeq Facility. Lastly, this work was supported by the Northwestern University Behavioral Phenotyping Core.

## Author contributions

ZC and SJS designed the project. ZC performed the experiments, analyzed the data, and assembled the data. ZC wrote the manuscript. CW, DG, and SJS revised the manuscript. KPF and HH contributed to the surgery and treatment injections. KPF, MBARI, and CW contributed to data acquisition of behavioral testing. MBARI, KPF, HW, and HH contributed to tissue processing. YW contributed to EdU assay and flow cytometry gating. AR contributed to library preparation for single-cell RNA coupled with TCR sequencing. WC and RB contributed to the analysis for single-cell RNA coupled with TCR sequencing and interpretation. ZC, KPF, MBARI, HH, VV, ZS, BTD, and SJS contributed to overall data interpretation. All authors reviewed the manuscript and approved its final submitted version.

## Data and code availability

All raw and processed data are available at: GEO pending release

All code can be found at: https://github.com/Jennieecc/aCD49d-study.git

