## Supplementary material for "antiCD49d Ab treatment ameliorates age-associated inflammatory response and mitigates CD8+ T-cell cytotoxicity after traumatic brain injury": supp file

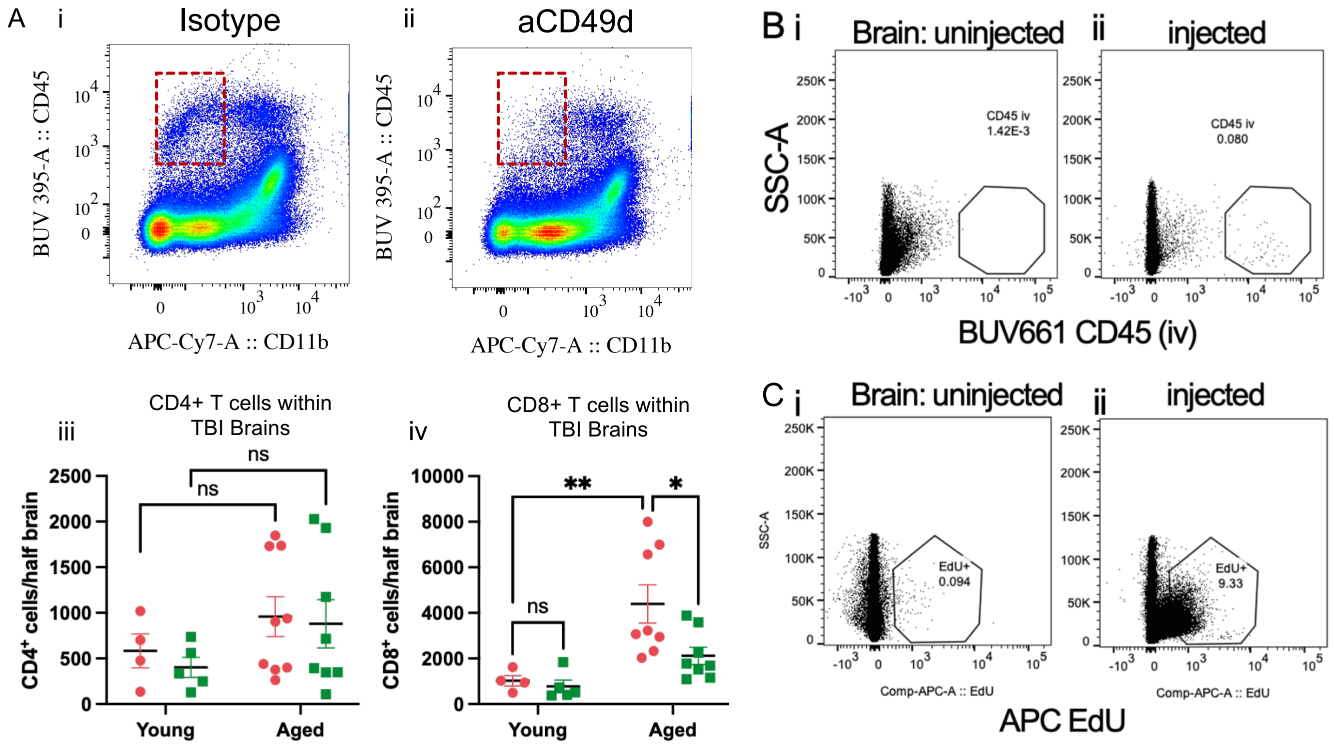


Supplement Figure 1. **aCD49d Ab treatment reduced CD8+ T cells in the aged brains and improved post-TBI deficits in motor function and working memory.** A. i-ii. Representative aCD49d Ab lymphocyte depletion in the aged brains are shown. iii-iv. aCD49d Ab treatment specifically reduced CD8+ T cells in the aged brains but not CD4+ T cells. B. i-ii. Representative BUV661 $CD45^+$ infiltrating cells in the brain are shown. C. i-ii. Representative $EdU^+$ proliferating cells in the brain are shown. Data are from from two independent experiments in A. All data are shown as the mean ± SEM, 2-way ANOVA with Tukey's multiple comparisons test. n = 5-9/group for A, *$p < 0.05$, **$p < 0.01$, ***$p < 0.001$.

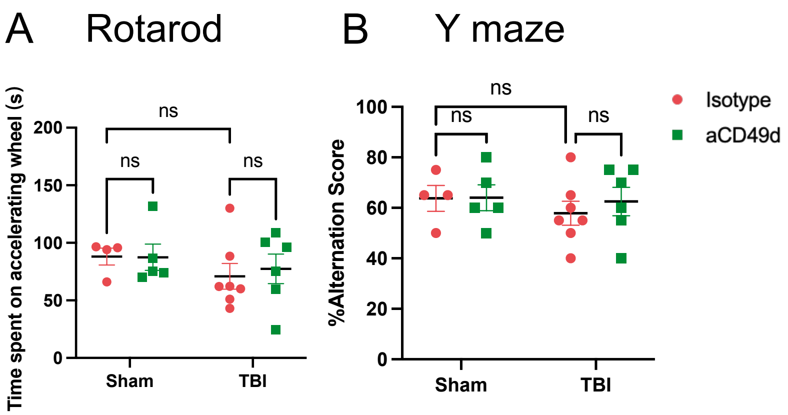


Supplement Figure 2. **No effect of aCD49d Ab was seen in the behavioral performances of young mice post TBI**. Results of A. rotarod indicated by time spent on accelerating wheels (s) and B. Y maze indicated by %alteration score. Data are from two independent experiments. All data are shown as the mean ± SEM, 2-way ANOVA with Tukey's multiple comparisons test. n = 8-10/group for A and B.

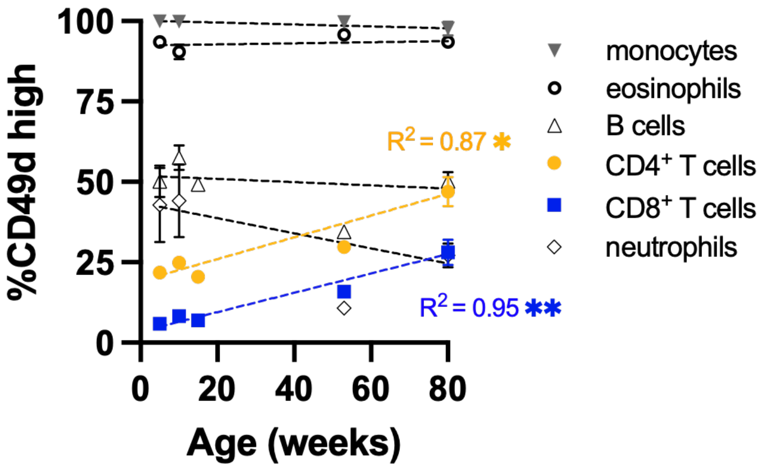


Supplement Figure 3. **Correlation of CD49d expression in different immune cells in the blood with age.** All data are shown as the mean ± SEM, Pearson's correlation analysis, n=15. *$p < 0.05$, **$p < 0.01$.

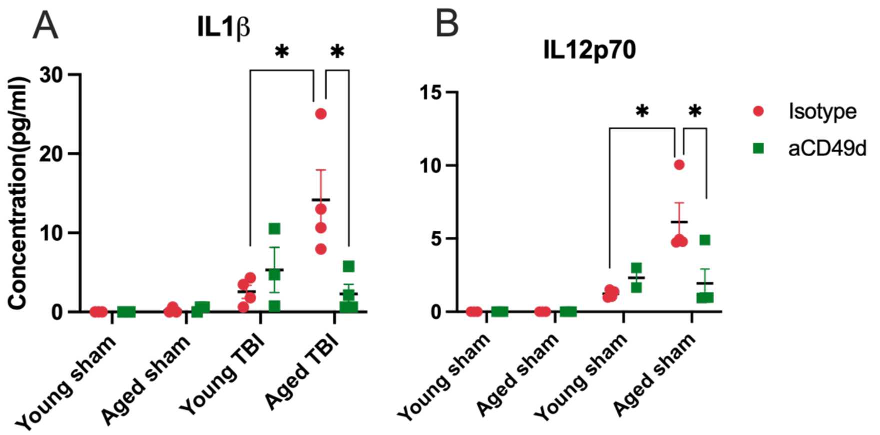


Supplement Figure 4. **Multiplex cytokine analysis in aged and young mice at 7 days post injury**. Levels of plasma cytokines including A. IL12p70 and B. IL1β. n =3-4/group, *p < 0.05. All data are from one independent experiment. Data are shown as the mean ± SEM, 2-way ANOVA with Tukey's multiple comparisons test.

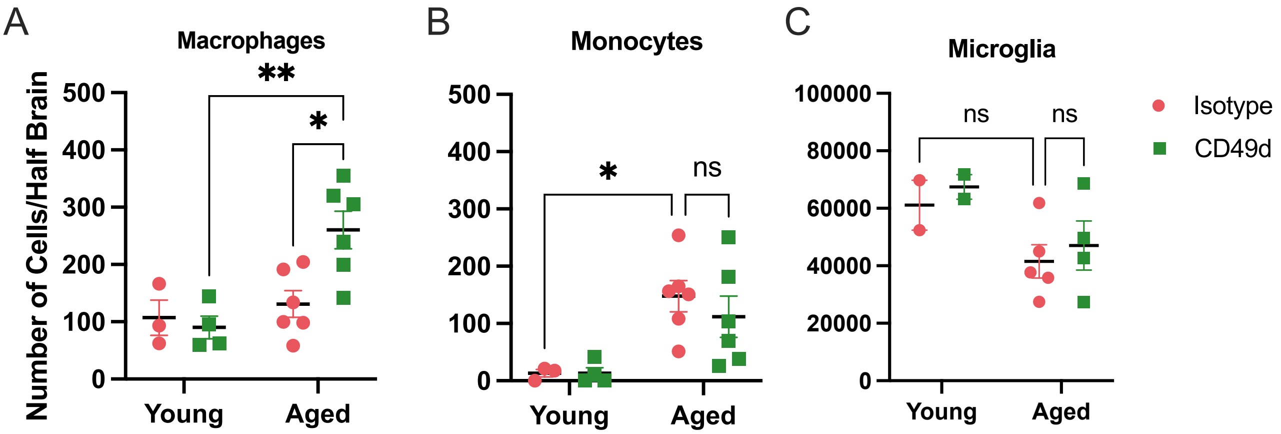


Supplement Figure 5. Quantifications of A. CD64+ macrophages, B. Ly6c+ monocytes and C. $CD45^{dim}CD11b+$ in both young and aged mouse brains 2 months post TBI. Data are from two independent experiments and shown as the mean ± SEM, 2-way ANOVA with Tukey's multiple comparisons test. n=2-6/group, *p < 0.05, **p < 0.01.

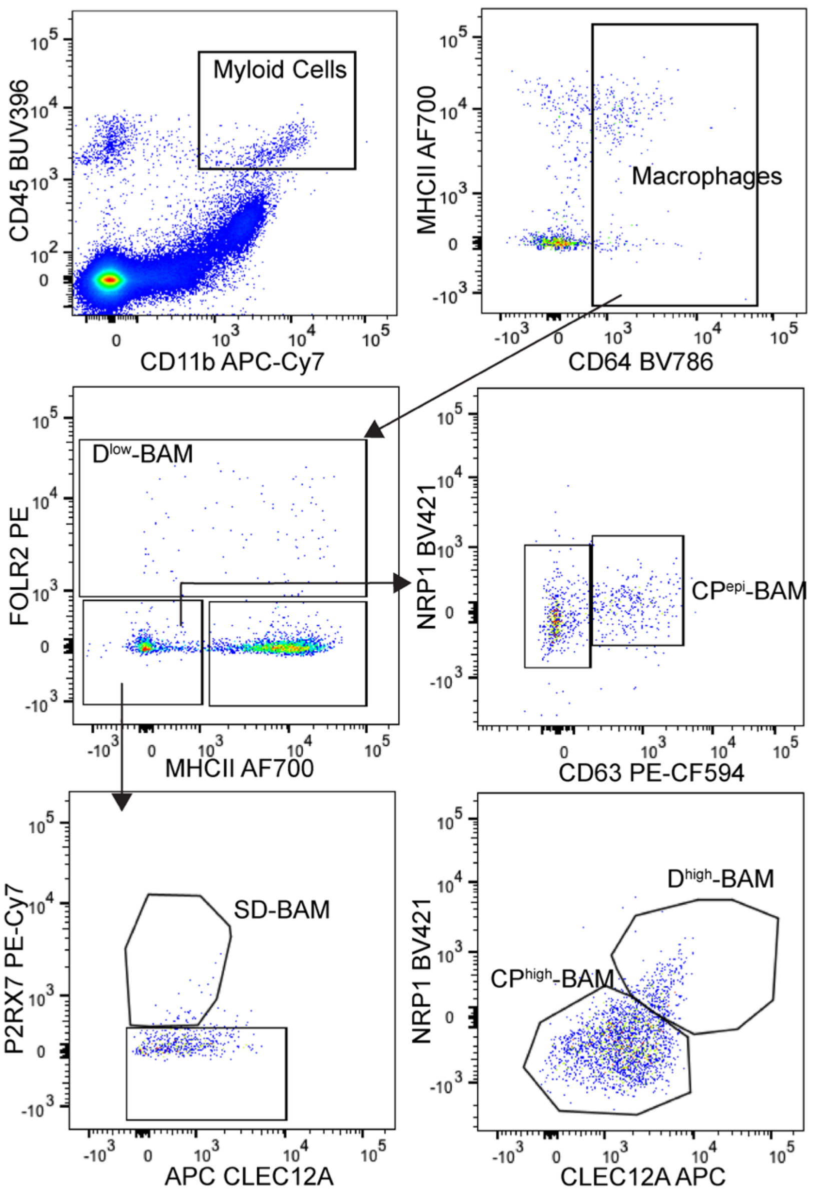


Supplement Figure 6. Gating strategy for tissue-specific brain resident macrophages or BAM. CD45+CD64+ macrophages were further gated on FOLR2 and NRP1 for dural BAM (D-BAM), MMR and P2RX7 for subdural BAM (SD-BAM), and CD63 for choroid plexus BAM (CP-BAM).

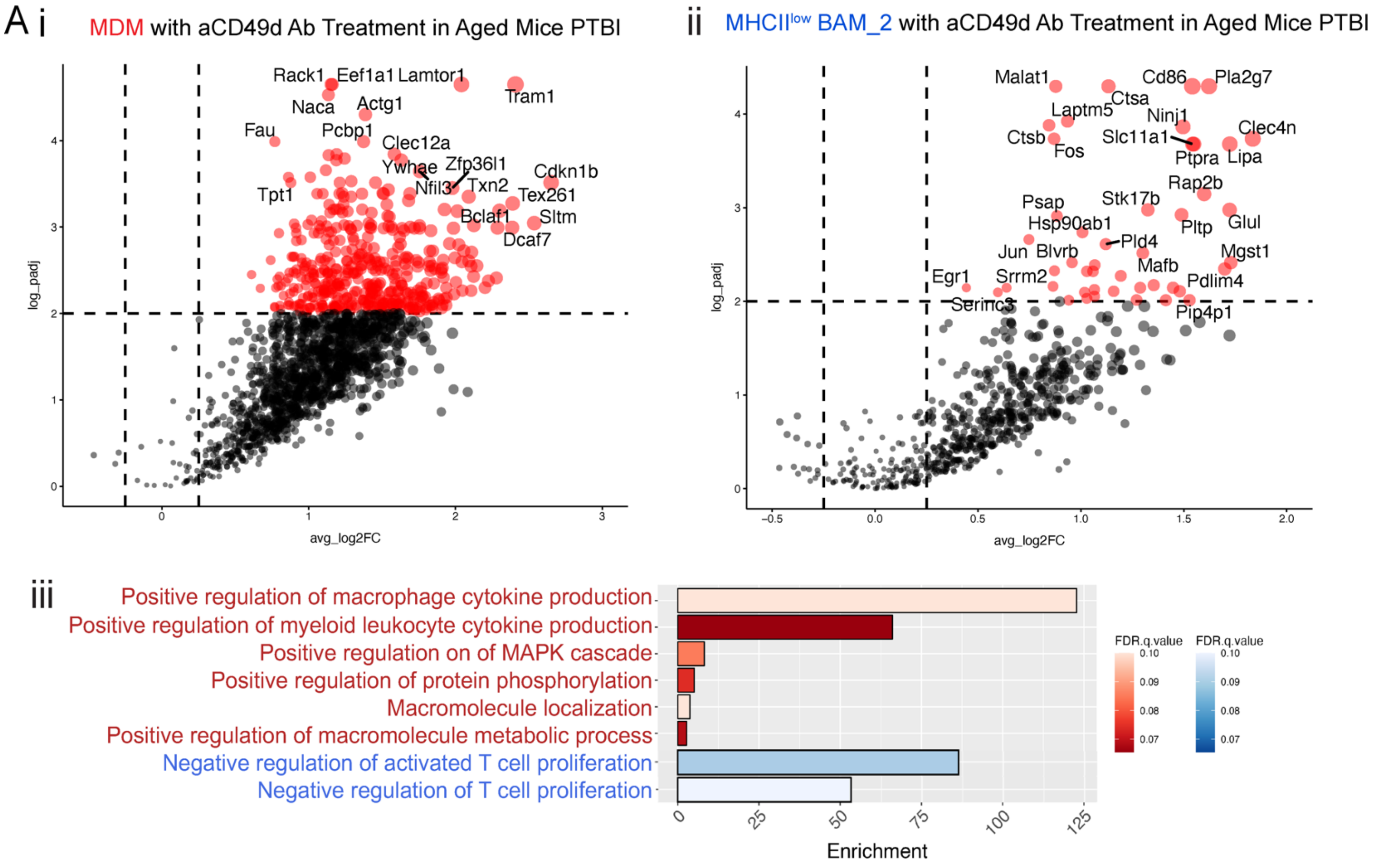


Supplement Figure 7. Volcano plots displaying genes that are DE (adjusted P<0.01, log2(FC)>0.25) between i. MDMs with aCD49d Ab versus isotype treatment and ii. BAMs with aCD49d Ab versus isotype treatment in aged mice two months post TBI. No downregulated DEs are found in both groups.

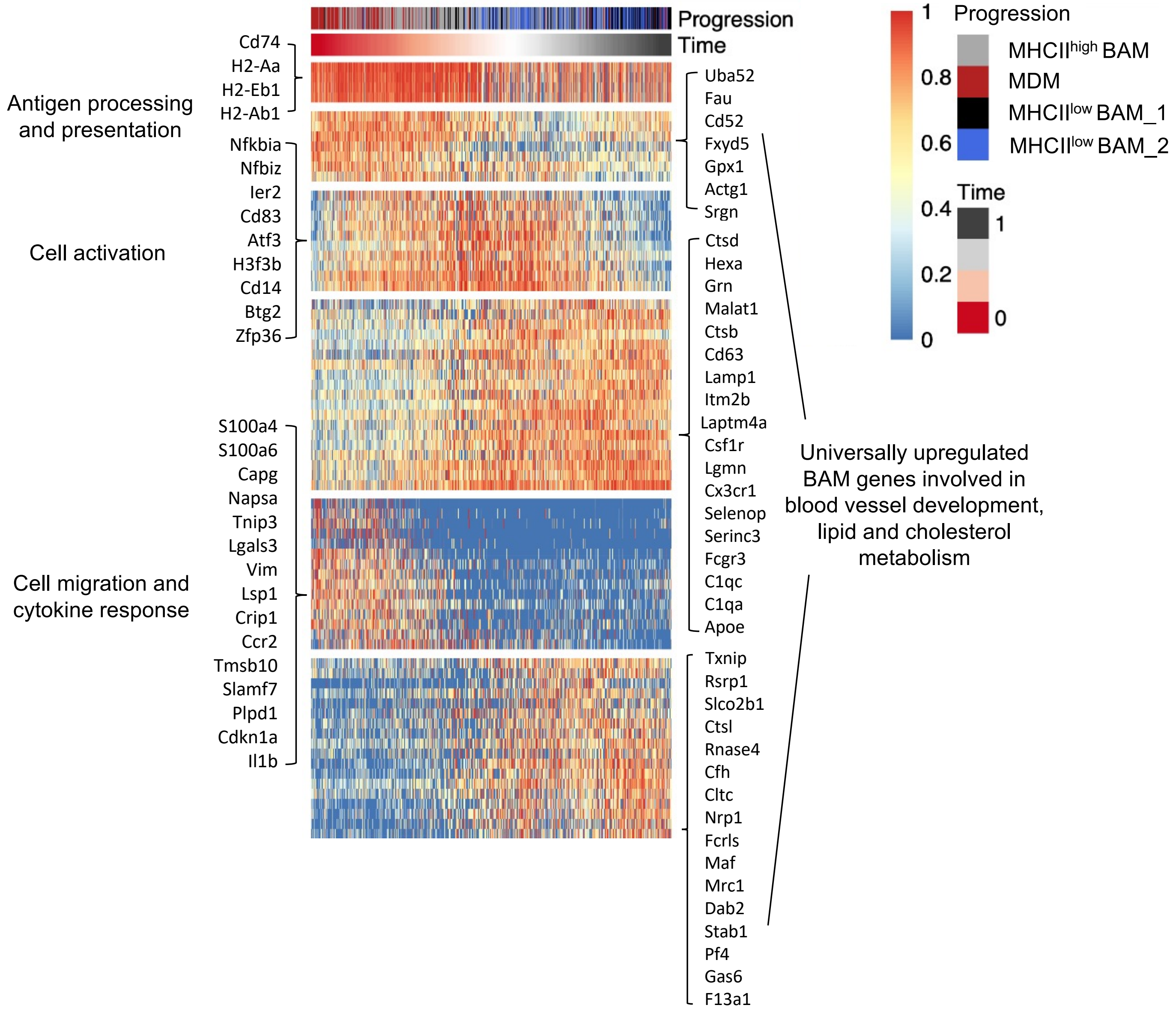


Supplement Figure 8. The top 100 genes were clustered into four modules that correspond to the cellular transition from MDM to BAMs.

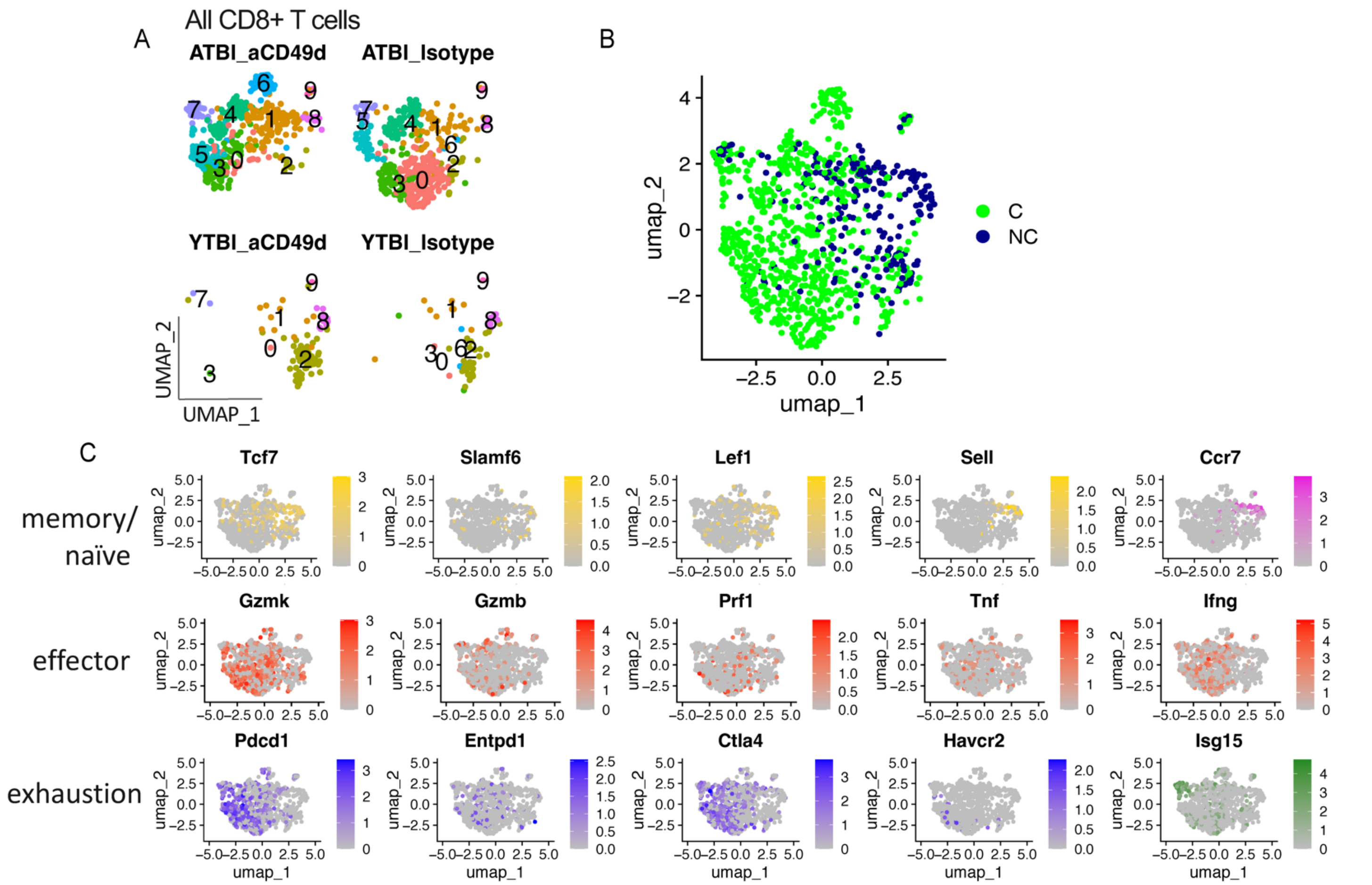


Supplement Figure 9. A. UMAP showing unsupervised clustering across samples. B. single-cell TCR analysis overlaid on UMAP projections showing distribution of CD8+ T cell clonality. C. Feature plots depicting various T cell differentiations states reveals that brain T cells from aged TBI mice had two pools of activated CD8+ T cells: dysfunctionally and functionally activated.

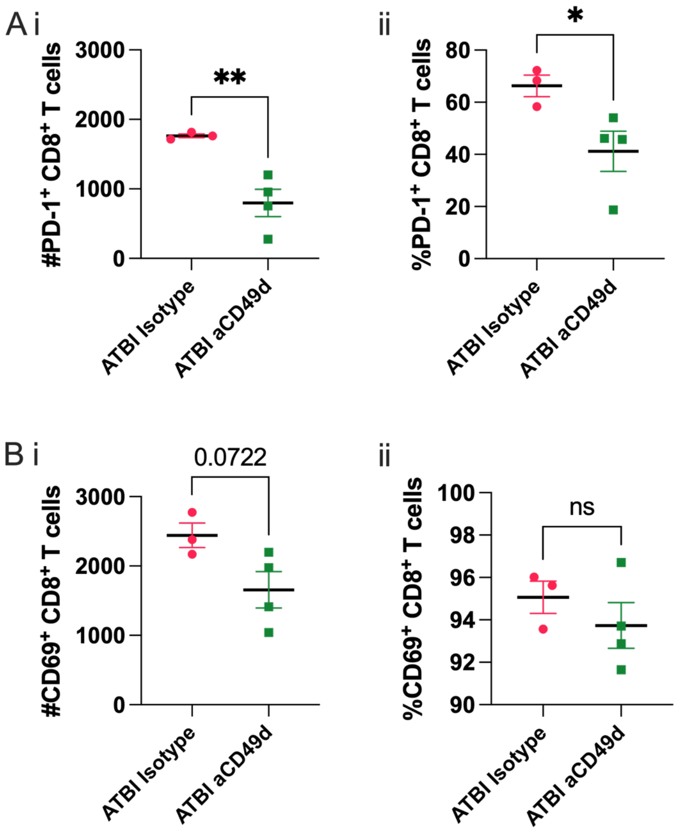


Supplement Figure 10. A. i Quantification of PD-1+ and ii. % PD-1+ CD8+T cells in all CD8+ T cells in aged mouse brains with isotype vs aCD49d Ab treatment. B. i Quantification of CD69+ and ii. CD69+ CD8 T+ cells in all CD8+ T cells in aged mouse brains with isotype vs aCD49d Ab treatment. n =3-4/group, *p < 0.05, **p<0.01. All data are from one independent experiment. Data are shown as the mean ± SEM, 2-way ANOVA with Tukey's multiple comparisons test.
